# Fine-mapping methods for complex traits: essential adaptations for samples of related individuals

**DOI:** 10.1101/2021.10.10.463846

**Authors:** Junjian Wang, Francesco Tiezzi, Yijian Huang, Julong Wei, Christian Maltecca, Jicai Jiang

**Affiliations:** Department of Animal Science, North Carolina State University, Raleigh, NC, 27695, USA; Department of Agriculture, Food, Environment and Forestry (DAGRI), University of Florence, Firenze, 50144, Italy; Smithfield Premium Genetics, Rose Hill, NC, 28458, USA; Center for Molecular Medicine and Genetics, Wayne State University, Detroit, MI, 48201, USA

**Keywords:** fine-mapping, complex trait, Bayesian, related individuals

## Abstract

Fine-mapping causal variants from GWAS loci is challenging in populations with substantial relatedness, such as livestock, as standard methods often assume unrelatedness, leading to poor fine-mapping accuracy. Here, we introduce a comprehensive Bayesian framework to address this. Our approach features BFMAP-SSS for individual-level data, which uses a linear mixed model (LMM) and shotgun stochastic search with simulated annealing. For summary statistics, we develop FINEMAP-adj and SuSiE-adj, novel strategies that directly use standard FINEMAP and SuSiE for samples of related individuals by employing LMM-derived inputs (particularly a relatedness-adjusted linkage disequilibrium matrix). Furthermore, gene-level posterior inclusion probability (PIP_gene_) is proposed to enhance detection power by aggregating variant signals. Extensive simulations based on pig genotypes show our methods substantially outperform existing tools (FINEMAP, SuSiE, FINEMAP-inf, SuSiE-inf, and GCTA-COJO) in samples of related individuals, achieving notable improvements in fine-mapping accuracy (e.g., up to several-fold increases in AUPRC). PIP_gene_ markedly improves candidate gene identification. Application to Duroc pig traits demonstrates practical utility. This work provides robust, validated methods and associated software and scripts for accurate fine-mapping in populations with complex relatedness.

## Introduction

Genome-wide association studies (GWAS) have successfully discovered many genetic associations with complex traits and diseases (1). However, these studies primarily identify associated genomic regions rather than pinpoint the actual causal variants, largely due to linkage disequilibrium (LD), whereby non-causal variants are correlated with true causal variants (2, 3). Such associated regions often contain many SNPs, the majority of which have no direct effect on the trait (4), thereby complicating downstream functional validation. Fine-mapping aims to address this by statistically prioritizing the most likely causal variants within a GWAS-identified locus, assuming at least one such variant exists (5) and thus narrowing the field for functional studies.

Statistical fine-mapping methods typically account for LD structure to identify a small set of putative causal variants, effectively treating the problem as variable selection in a regression model (6, 7). Given *m* SNPs in a region, 2^*m*^ possible causal models exist, making exhaustive evaluation computationally prohibitive for densely genotyped regions. Early approaches like conditional stepwise selection (e.g., GCTA-COJO) (8) identify independent signals but can be suboptimal, particularly when multiple causal variants in LD are present or when a non-causal variant strongly tags multiple causal variants (5, 7). Methods like CAVIAR (7) and its extension CAVIARBF (9) improved upon this by jointly modeling multiple variants and exhaustively evaluating all possible models with up to a pre-specified small number of causal variants, but faced computational limitations with larger regions or higher numbers of true causal variants. To address this challenge, newer approaches were developed. FINEMAP (10) employs shotgun stochastic search (SSS) (11) to efficiently explore the vast model space, while SuSiE uses Iterative Bayesian Stepwise Selection (IBSS) with the “Sum of Single Effects” model (12, 13) to search multiple effects. These methods have become widely adopted due to their improved computational efficiency. Recent extensions like FINEMAP-inf and SuSiE-inf (14) further refine these by attempting to model infinitesimal effects within the candidate region.

Despite these advancements, a major limitation of most existing fine-mapping methods, particularly those based on summary statistics, is their foundation in linear regression models that assume samples of unrelated individuals. The use of related individuals violates the underlying model assumption of unrelatedness, potentially distorting LD and subsequent causal variant identification. While it has become standard to use summary statistics derived from linear mixed models (LMMs) that account for relatedness and/or infinitesimal background at the GWAS stage (15, 16), the downstream fine-mapping tools often still do not correctly incorporate this relatedness structure into their own models. This critical issue of appropriately adapting fine-mapping for related individuals has particularly been overlooked in livestock genetics, where populations typically exhibit substantial and complex inter-individual relatedness due to breeding practices.

This study introduces a comprehensive Bayesian fine-mapping framework specifically designed to address these challenges in samples of related individuals. We aim to demonstrate that existing summary-statistics-based fine-mapping methods cannot be naively applied to samples of related individuals and develop comprehensive methodological solutions for related samples. We build upon our previously developed BFMAP approach, which uses an LMM for individual-level data and originally employed forward selection (BFMAP-Forward) (17). Here, we enhance the BFMAP framework by implementing a more robust model exploration strategy, BFMAP-Shotgun Stochastic Search (BFMAP-SSS), to better handle scenarios with highly correlated causal variants. Recognizing the widespread use of summary statistics, we further develop novel adaptation strategies, FINEMAP-adj and SuSiE-adj. These approaches enable the direct use of standard FINEMAP and SuSiE software with summary statistics obtained from related individuals by inputting LMM-derived inputs and a relatedness-adjusted LD matrix. Finally, to improve detection power in genomic regions with challenging LD patterns, as is common in livestock, we propose and validate the use of gene-level posterior inclusion probability (PIP_gene_) that aggregates variant-level evidence.

Through simulations based on real livestock genotypes and application to economically important traits in Duroc pigs, we demonstrate both the inadequacy of naively applying existing summary-statistics methods to related individuals and the substantial improvements offered by our comprehensive methodological solutions (BFMAP-SSS, FINEMAP-adj, SuSiE-adj, and PIPgene) in fine-mapping performance. This work provides both novel methodologies and practical adaptations essential for robust identification of causal variants and trait-associated genes in samples with complex genetic relatedness.

## Materials and methods

### Fine-mapping with unrelated individuals

For a sample of unrelated individuals, the genetic fine-mapping analysis can be represented using the following linear model:

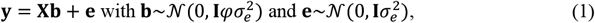

where **y** is an *n*-vector of covariate-adjusted phenotypes for *n* individuals, **X** is an *n*-by-*m* matrix of standardized genotypes for *n* individuals and *m* SNPs in a genomic region of interest, **b** is an *m*-vector of joint SNP effects, **e** is an *n*-vector of residual effects, and *φ* and 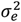 are hyperparameters related to variance components. Specifically, *φ* represents the ratio of per-SNP genetic variance to residual variance 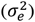, and we can set *φ*=0.01 to specify a moderate prior variance for SNP effects. Note that model (1) does not account for genetic relatedness between individuals, restricting its application to samples of unrelated individuals.

Model (1) can be expressed as follows:

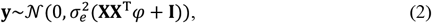

which is further transformed to 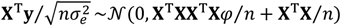. Denoting 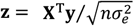 and **Σ** = **X**^T^**X**/*n* (we ignore the error introduced by the estimation of 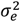 and the use of *n* instead of *n*−1), we have

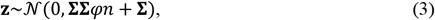

where **z** is an *m*-vector of z-scores from linear regression associations and **Σ** is an *m*-by-*m* matrix of Pearson’s correlations between *m* SNPs. It is important to note that model (3) holds for any set of SNPs in the genomic region. For a set of SNPs (denoted as *s*), let **z**_*s*_ and **Σ**_*s*_ represent the z-score vector and correlation matrix, respectively. Given model (3), we can compute the Bayes factor of the model (denoted as *M*) for *s* over the reduced model including no SNPs:

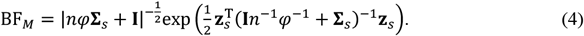

Equation (4) demonstrates the computation of Bayes factors for multi-SNP models using GWAS summary statistics and reference panel SNP correlations. This approach forms the foundation for genetic fine-mapping tools like CAVIARBF (9) and FINEMAP (10). Compared to individual-level data calculations under model (2), this method offers enhanced computational efficiency, particularly when leveraging reference panels like the 1000 Genomes Project (18) for SNP correlation estimation. However, like model (2), this approach is restricted to unrelated individuals.

Genetic fine-mapping is fundamentally a model selection problem. Although some fine-mapping methods, such as GCTA-COJO (8), use *P*-values rather than Bayes factors, they maintain consistency with Bayesian approaches. Specifically, when GCTA-COJO identifies a small conditional *P*-value for a SNP, this typically corresponds to a substantially larger log(*BF*) for the model including this SNP than for the comparable model without this SNP. Bayesian fine-mapping methods, such as CAVIARBF (9) and FINEMAP (10), evaluate sufficiently many multi-SNP models to compute posterior inclusion probability (PIP) for each variant in the candidate genomic region.

SuSiE (13) introduces a novel approach by decomposing the overall genetic effect into a sum of individual single effects, rather than explicitly evaluating different combinations of multiple SNPs. This computational innovation, combined with the implementation of variational Bayes for approximate posterior inference, enables SuSiE to identify multiple causal SNPs with high computational efficiency. Despite these methodological advances, SuSiE operates within the same linear framework as CAVIARBF and FINEMAP.

### Fine-mapping with related individuals: individual-level data

We previously developed a Bayesian framework for genetic fine-mapping with related individuals, which we refer to as BFMAP (17). BFMAP uses individual-level genotype and phenotype data based on the following linear mixed model:

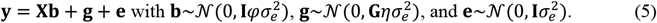

Model (5) extends model (1) by including an additional random-effect term, **g**, to account for genetic relatedness and whole-genome infinitesimal effects. **G** represents the genomic relationship matrix (GRM) (19), and *η* denotes the ratio of additive genetic variance to residual variance 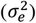. Other terms remain as defined in model (1).

For model implementation, we need to specify both *η* and 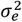. When the effects of candidate variants (**Xb**) account for only a small proportion of phenotypic variance (typically true when modeling variants from a small genomic region), we can set the value of *η* based on the SNP heritability (*h*^2^) by *η* = *h*^2^/(1−*h*^2^). In practice, we can use the SNP *h*^2^ estimate from the null model without the **Xb** term to determine *η*. For the residual variance 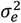, we assign the non-informative Jeffrey’s prior, 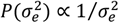.

For any model *M* that specifies a set of SNPs to be included, let *s* denote this SNP set. Given *D* = {**y, X, G**, *φ, η*}, we have the following equation for the marginal likelihood after integrating out 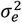:

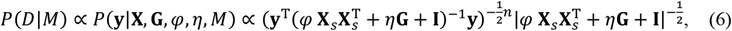

where **X**_*s*_ denotes the genotype matrix for SNP set *s*. The effective computation of *P*(*D*|*M*) has been detailed in our previous study (17).

Computing *P*(*D*|*M*) for every possible model is computationally infeasible. We have implemented two methods in BFMAP to facilitate model selection: one based on forward selection and the other using shotgun stochastic search (SSS) (11), which we refer to as BFMAP-Forward and BFMAP-SSS, respectively. The implementation of BFMAP-Forward has been detailed in our previous study (17). The forward selection procedure sequentially adds independent signals to the model, repositions them for refinement, and identifies variants related to each signal. For BFMAP-SSS, we largely follow the implementation in FINEMAP (10). SSS efficiently explores the vast model space by evaluating neighboring models at each iteration. However, a naïve implementation may fail to discover causal variants even with a very long chain for certain LD structures. To address this limitation, we incorporate simulated annealing (20) to ensure more thorough search. We apply the following linear cooling scheme:

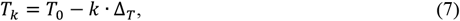

where *T*_0_ is the initial temperature, *T*_*k*_ is the temperature at the *k*-th step, and Δ_*T*_ is the temperature decrement per step. For all analyses in this study, *T*_0_ was set to 100, and Δ_*T*_ was fixed at 1. We performed 100 SSS iterations at each temperature step and set the final temperature to 1. An additional 100 SSS iterations were conducted at this final temperature to ensure thorough exploration of the model space.

### Fine-mapping with related individuals: summary statistics

In this section, we present a summary-statistics-based fine-mapping method for related individuals, following the framework of BFMAP (17). Denoting **H** = **G***η* + **I**, we transform model (5) to:

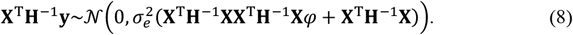

Defining a diagonal matrix **D** = diag(**X**^T^**H**^−1^**X**) and an *m*-by-*m* matrix 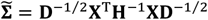, we obtain from (8):

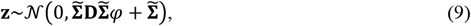

where 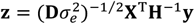 (that is, an *m*-vector of z-scores from linear mixed model associations). For a set of SNPs (denoted as *s*) in a candidate genomic region, we can readily derive the Bayes factor of the model (denoted as *M*) for *s* over the reduced model including no SNPs:

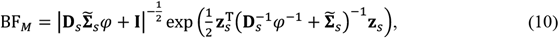

where **z**_*s*_, **D**_*s*_, and 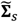 represent **z, D**, and 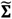, respectively, corresponding to SNP set *s*.

It is important to clarify the treatment of 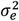 in equation (10). While our BFMAP framework (e.g., equation (6)) involves integrating out 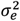 using its Jeffrey’s prior, the z-scores used as input for equation (10) inherently treat 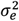 as a fixed quantity. This is because z-scores are standardized statistics that already incorporate an estimate of 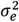. However, this difference in treatment is generally of little practical consequence. For reasonably large samples (*n* > 1,000), the estimate of 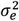 is highly precise. Therefore, using z-scores that embed a point estimate of 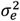 typically yields results very similar to those that would be obtained from a full Bayesian integration of 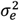.

Equation (10) has a similar form to equation (4). Specifically, while the genotype correlation matrix in equation (4) for unrelated individuals is defined as **Σ** = **X**^′^**X**/*n*, this is replaced in equation (10) for related individuals by 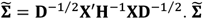 can be regarded as relatedness-adjusted genotype correlations, which may substantially differ from **Σ**. Using 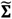 instead of **Σ** is critical for fine-mapping with related individuals. Misuse of unadjusted correlations can lead to poor performance.

Applying the approximation **D** ≈ **I***ñ* (where *ñ* is the average of the diagonal elements of **D** and **I** is the identity matrix) to equation (10) transforms it into equation (11):

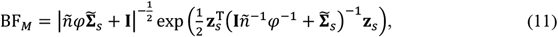

which has a structural form the same as equation (4). This approximation mirrors that used by GRAMMAR-Gamma (21), 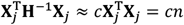, where **X**_*j*_ represents the vector of standardized genotypes for SNP *j* and *c* is a constant across all SNPs. The structural similarity between equations (11) and (4) enables direct use of existing summary-statistics-based fine-mapping tools (e.g., FINEMAP and SuSiE) for related individuals. The use involves two steps: 1) calculate **z**, *ñ*, and 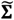 as described in this section, and 2) input these as z-scores, sample size, and LD matrix, respectively, into FINEMAP and SuSiE. We refer to these adapted methods as FINEMAP-adj and SuSiE-adj.

### Posterior probability metrics

Given the Bayes factor BF_*M*_ or the marginal likelihood *P*(*D*|*M*) of model *M* (which defines a specific set of candidate causal SNPs, *s*), its posterior model probability is calculated as:

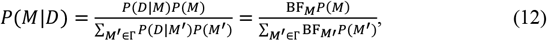

where Г represents the model space and *M*^′^ represents any individual model in the model space. We use a standard binomial prior, *P*(*M*) = *π*^|*M*|^(1 − *π*)^*m*−|*M*|^, to induce sparsity in BFMAP, where |*M*| is the model size (number of SNPs in *M*) and *π* is the prior probability that a SNP is causal. This prior implies an expected model size of *mπ*. Following standard fine-mapping practice (9, 10), *π* is set to 1/*m*, reflecting an expectation of one causal variant *a priori*.

The posterior inclusion probability (PIP) quantifies the probability that a specific SNP has a non-zero effect, given the observed data and priors (22). For SNP *j*, the PIP is computed as:

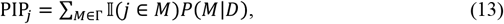

where 𝕀(*j* ∈ *M*) is an indicator function returning 1 if *j* is included in model *M* and 0 otherwise. This definition of PIP is used by methods that explore a broad model space, such as FINEMAP, SuSiE, FINEMAP-inf, SuSiE-inf, FINEMAP-adj, SuSiE-adj, and BFMAP-SSS.

Our BFMAP-Forward implementation, however, operates via a stepwise forward selection procedure. Consequently, it calculates a posterior conditional inclusion probability (PCIP). A variant’s PCIP is the posterior probability of the variant having a non-zero effect, conditional on the set of variants already incorporated into the model. While PCIPs are by definition conditional (unlike marginal PIPs), they serve as the direct analogue to PIPs for BFMAP-Forward throughout this study.

Finally, to aggregate evidence at the gene level, we propose gene-level PIP (PIP_gene_), defined as the posterior probability that a given gene harbors at least one non-zero-effect variant. This metric is computed by summing the posterior probabilities of all models containing at least one SNP within the gene:

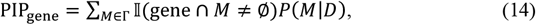

where 𝕀(gene ∩ *M* ≠ ∅) indicates whether any SNP in the gene is included in model *M*. This gene-level aggregation offers a complementary perspective involving a trade-off between localization precision and power. For PIP_gene_ calculation, gene boundaries were defined using Ensembl annotations and extended by 3kb upstream and downstream to include potential regulatory regions.

### Data simulation

We evaluated fine-mapping methods using genotype data from 31,301 Duroc pigs provided by Smithfield Premium Genetics (Roanoke Rapids, NC, USA). The animals were genotyped with the PorcineSNP60 BeadChip (Illumina Inc., San Diego, CA, USA) and genotypes were imputed to 34,615,361 autosomal variants using SWIM 1.0 (23). This full set of imputed variants was then filtered for minor allele frequency (MAF) ≥1%, Hardy-Weinberg equilibrium (*P* ≥1 × 10^−8^), and IMPUTE2 info score ≥0.8, resulting in ∼11.7 million quality-controlled variants on autosomes. From these quality-controlled variants, we randomly selected a 4-Mb region on chromosome 1 containing 6,504 variants for the simulation study.

Quantitative traits were simulated using model (5) with a heritability of 0.5 using MPH (24). The GRM **G**, used to model the covariance of polygenic effects due to genetic relatedness, was computed using 10,000 randomly selected, quality-controlled chip SNPs. We positioned 1, 2, or 3 causal variants within the central 2-Mb region, collectively explaining either 1% or 4% of phenotypic variation (PVE). Simulations used samples of 5,000 or 10,000 randomly selected individuals, generating 100 replicates for each of the 12 scenarios (3 causal variant counts × 2 PVE levels × 2 sample sizes).

For gene-based fine-mapping evaluation, we partitioned the 4-Mb candidate region into 100 equal blocks to serve as gene proxies. This block size approximates the average pig gene length of ∼33 kb derived from the Sscrofa11.1 genome annotation (Ensembl release 113) (25). To generate the required summary statistics for fine-mapping from each simulated dataset, we conducted association tests for variants in the candidate region using two approaches: 1) a linear mixed model (LMM) implemented in GCTA-MLMA (v1.94.1) (26), and 2) a linear regression model in PLINK 2.0 (−-glm) (27) using the top 20 genotype principal components as covariates for population structure control. GCTA-MLMA used the same GRM as in the trait simulation. Furthermore, required LD matrices were generated from each simulation sample’s genotype data: both the standard SNP correlation matrix (**Σ**) as in model (3) and the relatedness-adjusted correlation matrix 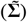 as in model (9). The standard matrix (**Σ**) was used by GCTA-COJO, FINEMAP, SuSiE, FINEMAP-inf, and SuSiE-inf, while the relatedness-adjusted matrix 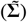 was used by FINEMAP-adj and SuSiE-adj.

### Performance evaluation in simulations

The performance of various fine-mapping approaches was benchmarked using the datasets generated across the 12 distinct simulation scenarios (3 causal variant counts × 2 PVE levels × 2 sample sizes). It is important to note that while 100 replicates were generated for each scenario, not all replicates resulted in a significant association signal within the candidate region when analyzed by GCTA-MLMA. Specifically, only those simulation replicates where at least one variant in the candidate region achieved a GCTA-MLMA *P*-value < 1 × 10^−5^ were retained for subsequent fine-mapping evaluation. This yielded a minimum of 74 such replicates for any given scenario, with detailed counts available in Supplementary Table S1.

The methods evaluated included our novel approaches (BFMAP-SSS, FINEMAP-adj, SuSiE-adj, and PIP_gene_), our previously described BFMAP-Forward method (17), and several well-established reference methods: FINEMAP v1.4.2 (10), SuSiE (as implemented in susieR v0.12.35) (12, 13), FINEMAP-inf v1.3, SuSiE-inf v1.4 (14), and GCTA-COJO (−-cojo-slct in GCTA v1.94.1) (8, 28). BFMAP-Forward and BFMAP-SSS were implemented in BFMAP v0.65, while FINEMAP-adj and SuSiE-adj directly used FINEMAP and SuSiE, respectively, with adapted inputs designed for related individuals. For consistency across all applicable methods, a causal variant’s prior PVE and the maximum number of causal variants were set to 0.01 and 5, respectively, in all simulation datasets. Furthermore, the same GRM, constructed from the 10,000 quality-controlled chip SNPs as detailed in the trait simulation, was used by all our LMM-based fine-mapping methods requiring such an input, specifically BFMAP (both BFMAP-Forward and BFMAP-SSS) and our -adj approaches (FINEMAP-adj and SuSiE-adj). For a comprehensive comparison, reference methods were applied twice, using association statistics as input from LMM (GCTA-MLMA) and, separately, from linear regression (PLINK 2.0 --glm).

We applied consistent *P*-value-based pre-filtering for fine-mapping across methods. Specifically, for GCTA-COJO (−-cojo-slct), the significance threshold in its stepwise procedure (−-cojo-p) was set to 1 × 10^−3^. Similarly, for all other fine-mapping tools evaluated (FINEMAP, SuSiE, FINEMAP-inf, SuSiE-inf, and our BFMAP and -adj approaches), only variants with a GWAS *P*-value < 1 × 10^−3^ from the respective input summary statistics (GCTA-MLMA or PLINK 2.0 --glm) were included in the fine-mapping analysis. This initial filtering threshold was chosen to reduce the computational burden of model searching for each tool while aiming to be sufficiently inclusive of potentially causal variants. Unless otherwise specified, all other parameters for these software tools were maintained at their default values.

To evaluate fine-mapping accuracy, variants from each method were designated as putatively causal based on their respective PIPs (or PCIPs for BFMAP-Forward) using thresholds ranging from 0.0 to 1.0. Performance was primarily quantified using recall (also known as sensitivity or power) and precision (also known as positive predictive value). Recall is the proportion of true causal variants correctly identified (calculated as TP / (TP + FN)), and precision is the proportion of designated putatively causal variants that are indeed true causal (calculated as TP / (TP + FP)). Here, TP, FP, and FN represent the number of true positives, false positives, and false negatives, respectively, determined at a given PIP (or PCIP) threshold and averaged over all simulation replicates retained for fine-mapping evaluation per scenario (Supplementary Table S1). Precision and recall were also calculated for gene-level PIPs using the same approach, with genes rather than variants as the unit of evaluation.

Given that GCTA-COJO (−-cojo-slct) does not compute PIPs, a modified evaluation strategy was necessary. Variants identified by GCTA-COJO were ranked by first prioritizing those from its jma output, followed by variants in its cma output, with the latter ranked by their conditional *P*-values. This *P*-value-based ranking was then used to assess GCTA-COJO’s ability to prioritize true causal variants relative to the PIP/PCIP-based rankings from other methods. Specifically, we evaluated its recall by selecting the top *k*-ranked variants (where *k* ranged from 1 to 20) as putatively causal. For each *k*, recall was calculated as the proportion of all true causal variants that were included in this top *k* selection, averaged across all qualifying simulation replicates per scenario.

### Duroc pig trait analysis

This study also analyzed several economically important traits from the same cohort of Duroc pigs described in the ‘Data simulation’ section. Phenotype data, collected by Smithfield Premium Genetics between 2015 and 2021, included growth traits for ∼27,000 animals (off-test body weight [WT], back fat thickness [BF], and loin muscle depth [MS]) and reproduction traits for 3,290 sows (number of piglets born alive [NBA], born dead [NBD], and weaned [NW]). The three growth traits were collected at the end of the performance test, as detailed in Bergamaschi et al. (29). All phenotypes were pre-adjusted to remove systematic environmental effects, including farm and physiological sources of variation. For association and fine-mapping analyses, we utilized the ∼11.7 million quality-controlled autosomal variants (derived as detailed in the ‘Data simulation’ section).

GWAS was performed using SLEMM-GWA v0.89.5 (30), which provides accuracy comparable to GCTA-MLMA but with orders of magnitude greater computational efficiency (Supplementary Figure S1). Association peaks were identified by visual inspection of the Manhattan plot for each trait, with a *P* -value threshold of 1 × 10^−5^. For each retained peak, candidate region boundaries were established by extending 2 Mb upstream and downstream from the position of the minimum *P*-value variant. Variants with *P* <1 × 10^−3^ within each defined candidate region were then retained for subsequent fine-mapping analyses. The same 10,000 quality-controlled chip SNPs selected for the trait simulation were used to construct the GRM for methods requiring it (e.g., SLEMM and our BFMAP and -adj approaches).

## Results

### Method overview

This study introduces a framework for fine-mapping complex traits in samples of related individuals, addressing analyses with both individual-level data and summary statistics. Figure 1 provides a schematic representation of this framework, illustrating the relationships between existing methods and the novel approaches developed herein.

**Figure 1.**
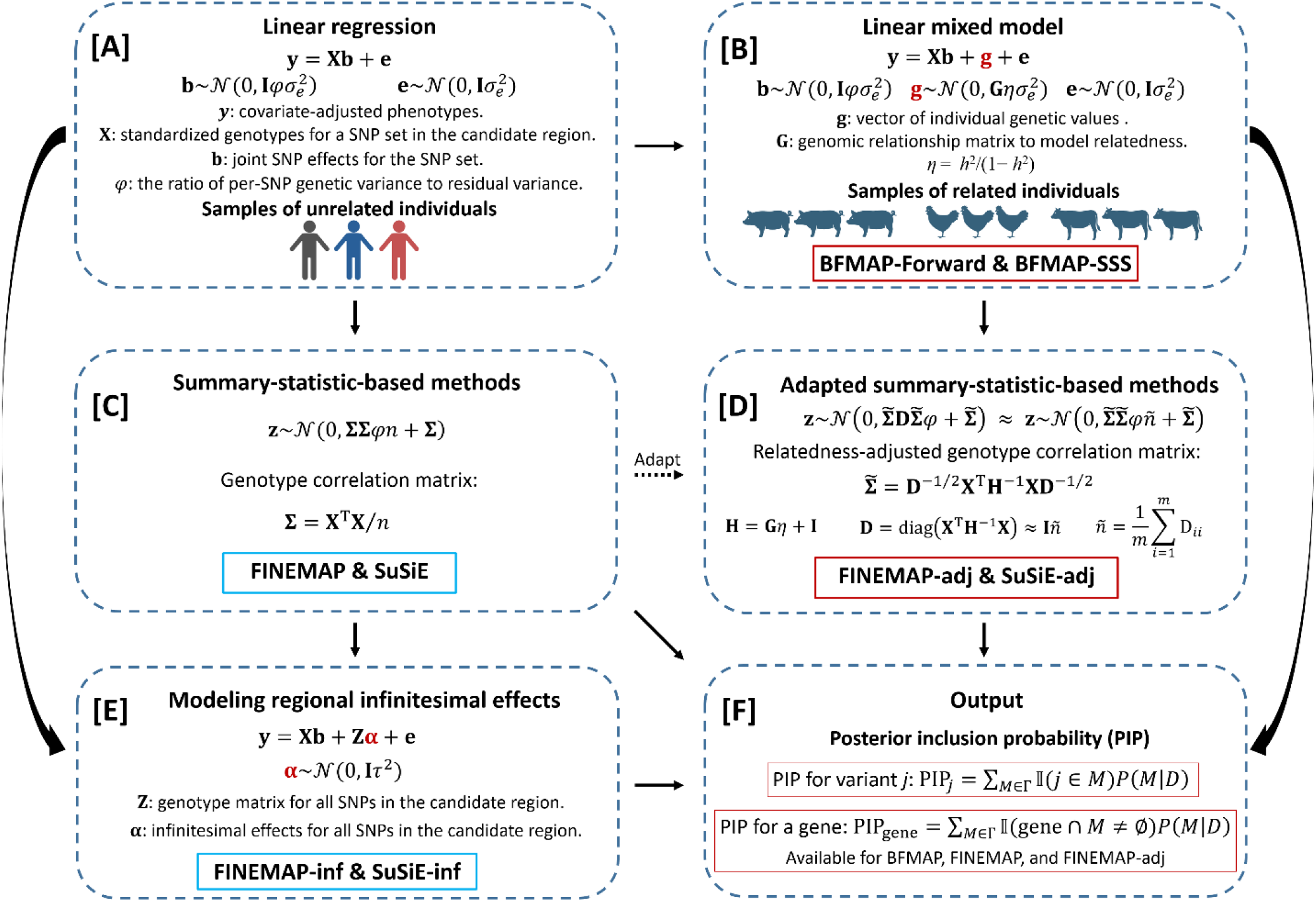
Overview of fine-mapping methods for unrelated and related individuals. Overview of fine-mapping approaches for samples of unrelated (A, C, and E) and related (B, D, and F) individuals. (A) Linear regression provides the foundation for fine-mapping in samples of unrelated individuals. (B) BFMAP employs linear mixed models with genomic relationship matrix for fine-mapping in samples of related individuals. (C) Standard summary-statistics methods (FINEMAP and SuSiE) are derived from the linear regression framework. (D) Adapted methods (FINEMAP-adj and SuSiE-adj) use relatedness-adjusted correlations 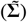 and effective sample size (*ñ*) derived from the BFMAP framework. Through the approximation **D** ≈ **I***ñ*, these methods achieve compatibility with standard FINEMAP and SuSiE implementations. (E) FINEMAP-inf and SuSiE-inf extend their respective base methods by modeling infinitesimal effects within candidate regions. (F) All methods produce variant-level posterior inclusion probability (PIP; or PCIP for BFMAP-Forward) as output. Gene-level PIP, available for methods that provide posterior model probabilities (BFMAP-Forward, BFMAP-SSS, FINEMAP-adj, and FINEMAP), aggregates variant-level evidence for prioritizing causal genes.

Typical fine-mapping tools, such as FINEMAP and SuSiE, usually use summary statistics. Their underlying models are derived from linear regression assuming samples of unrelated individuals, which limits their direct applicability or accuracy when substantial relatedness is present (Figures 1A and 1C). To overcome this, we developed BFMAP, a Bayesian fine-mapping approach first introduced for individual-level livestock data using forward selection (BFMAP-Forward) (17). BFMAP employs a linear mixed model (LMM) that explicitly accounts for whole-genome infinitesimal effects and genetic relatedness through a genomic relationship matrix (GRM) (Figure 1B). In this study, alongside the established BFMAP-Forward, we introduce BFMAP-SSS, a novel BFMAP implementation coupling shotgun stochastic search (SSS) with simulated annealing for enhanced model space exploration.

Building upon the BFMAP framework, we developed a strategy to leverage its advantages within a summary statistics context. The BFMAP model, initially designed for individual-level data, can be transformed into an equivalent model based on summary statistics (Figure 1D). Through an approximation (**D** ≈ **I***ñ*), analogous to the one used in GRAMMAR-Gamma, this transformed model yields a functional form identical to that underlying standard FINEMAP and SuSiE. This key structural equivalence enables the direct adaptation of these existing tools for accurate fine-mapping in related individuals. This adaptation involves a two-step process: first, calculating relatedness-aware inputs (including standard LMM-derived z-scores (**z**), an effective sample size (*ñ*), and a relatedness-adjusted linkage disequilibrium (LD) matrix 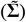, as detailed in Materials and methods); and second, providing these as the conventional z-score, sample size, and LD matrix inputs to FINEMAP or SuSiE. We term these adapted methods FINEMAP-adj and SuSiE-adj.

It is noteworthy that recently proposed fine-mapping methods, FINEMAP-inf and SuSiE-inf, are conceptually related to our LMM-based approaches (BFMAP, FINEMAP-adj, and SuSiE-adj) in that both incorporate terms to account for infinitesimal effects (Figure 1E). However, a key distinction is that FINEMAP-inf and SuSiE-inf model the infinitesimal effects of variants within the candidate fine-mapping region, whereas our methods model whole-genome infinitesimal effects via GRM.

Finally, recognizing the challenges of fine-mapping to single-variant resolution in populations with extensive LD, such as livestock, we propose the use of gene-level posterior inclusion probability (PIP_gene_). This metric aggregates variant-level evidence to assess the overall evidence for a gene harboring at least one non-zero-effect variant, offering a balance between localization precision and detection power (Figure 1F). In the subsequent sections, the performance of our novel contributions (BFMAP-SSS, FINEMAP-adj, and SuSiE-adj), alongside our previously developed BFMAP-Forward approach (17), is evaluated in comparison with existing approaches (FINEMAP, SuSiE, FINEMAP-inf, SuSiE-inf, and GCTA-COJO) for fine-mapping in related individuals.

### Simulation benchmarks

To rigorously evaluate the fine-mapping approaches, we conducted simulation studies based on real Duroc pig genotypes across 12 distinct scenarios. These scenarios varied by the number of true causal variants (1, 2, or 3), the total proportion of variance explained (PVE) by these causal variants (1% or 4%), and the sample size (5,000 or 10,000 individuals). For each scenario, 100 replicates were generated, though only those yielding significant GWAS associations (GCTA-MLMA *P* < 1 × 10^−5^) were retained for fine-mapping evaluation, ensuring robust downstream analysis (see Materials and methods for details).

#### Impact of relatedness adjustment on genotype correlations

Summary-statistics-based fine-mapping methods critically rely on an accurate LD matrix reflecting correlations between genotypes. In samples of related individuals, one might naively compute a standard LD matrix directly from the observed genotypes. However, as we have theoretically established in Material and methods (Section ‘Fine-mapping with related individuals: summary statistics’), a relatedness-adjusted LD matrix 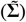 is required to properly account for the genetic relatedness among individuals in fine-mapping. To empirically illustrate the impact of this adjustment, we compared elements of the standard LD matrix (**Σ**) with those of the relatedness-adjusted LD matrix 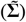 within our 4-Mb candidate region for simulations (Supplementary Figure S2). This comparison revealed substantial differences between the two types of correlations. Notably, the relatedness adjustment (transforming **Σ** to 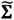) often led to a shrinkage of correlation magnitudes, particularly for variants that exhibited strong positive or negative correlations in the standard **Σ** matrix. Consequently, applying fine-mapping methods with a standard, unadjusted LD matrix in samples of related individuals, rather than the appropriate relatedness-adjusted version, can severely compromise fine-mapping performance and the accuracy of causal variant identification.

#### Performance of reference methods with different input association statistics

We first assessed the performance of established reference fine-mapping methods (FINEMAP, SuSiE, FINEMAP-inf, SuSiE-inf, and GCTA-COJO) using two types of input association statistics: those derived from a LMM (via GCTA-MLMA) accounting for relatedness and whole-genome infinitesimal effects, and those from a standard linear regression model (via PLINK 2.0 --glm) with top 20 genotype principal components for population structure control.

GCTA-COJO consistently demonstrated slightly superior performance when using LMM-derived association statistics across all simulation scenarios (Supplementary Figure S3). For the other reference methods (FINEMAP, SuSiE, and their -inf variants), the choice of input statistics had a more nuanced impact. In scenarios with 1 or 2 causal variants, these methods generally favored linear regression inputs in 4% PVE settings, whereas LMM-derived statistics were often preferred in 1% PVE settings, as reflected by precision-recall curves (Figure 2) and area under the precision-recall curve (AUPRC) values (Table 1). This distinction was less apparent in scenarios with 3 causal variants. Notably, SuSiE-inf frequently encountered convergence issues when using LMM-derived association statistics but performed robustly with linear regression inputs.

**Table 1.** AUPRC for causal variant identification by fine-mapping methods across simulation scenarios.

| Sample size | Method | Number of causal variants |  |  |  |  |  | Average |
| --- | --- | --- | --- | --- | --- | --- | --- | --- |
|  |  | 1 |  | 2 |  | 3 |  |  |
|  |  | PVE=0.01 | PVE=0.04 | PVE=0.01 | PVE=0.04 | PVE=0.01 | PVE=0.04 |  |
| 5,000 | BFMAP-SSS | 0.128 | 0.419 | 0.052 | 0.223 | 0.028 | 0.093 | 0.157 |
|  | BFMAP-Forward | 0.124 | 0.379 | 0.032 | 0.143 | 0.011 | 0.061 | 0.125 |
|  | FINEMAP-adj | 0.126 | 0.381 | 0.051 | 0.173 | 0.030 | 0.080 | 0.140 |
|  | SuSiE-adj | 0.117 | 0.388 | 0.036 | 0.159 | 0.026 | 0.066 | 0.132 |
|  | FINEMAP (LMM) | 0.067 | 0.064 | 0.024 | 0.020 | 0.013 | 0.013 | 0.034 |
|  | FINEMAP (LM) | 0.033 | 0.169 | 0.010 | 0.042 | 0.006 | 0.020 | 0.047 |
|  | FINEMAP-inf (LMM) | 0.033 | 0.021 | 0.016 | 0.012 | 0.010 | 0.011 | 0.017 |
|  | FINEMAP-inf (LM) | 0.026 | 0.139 | 0.011 | 0.044 | 0.004 | 0.023 | 0.041 |
|  | SuSiE (LMM) | 0.120 | 0.192 | 0.032 | 0.032 | 0.019 | 0.022 | 0.070 |
|  | SuSiE (LM) | 0.037 | 0.187 | 0.009 | 0.039 | 0.005 | 0.019 | 0.049 |
|  | SuSiE-inf (LMM) | 0.021 | 0.027 | 0.009 | 0.016 | 0.005 | 0.009 | 0.015 |
|  | SuSiE-inf (LM) | 0.019 | 0.201 | 0.008 | 0.040 | 0.003 | 0.018 | 0.048 |
| 10,000 | BFMAP-SSS | 0.188 | 0.468 | 0.096 | 0.378 | 0.077 | 0.199 | 0.234 |
|  | BFMAP-Forward | 0.140 | 0.469 | 0.053 | 0.250 | 0.061 | 0.157 | 0.188 |
|  | FINEMAP-adj | 0.189 | 0.463 | 0.089 | 0.303 | 0.076 | 0.188 | 0.218 |
|  | SuSiE-adj | 0.207 | 0.478 | 0.086 | 0.225 | 0.066 | 0.149 | 0.202 |
|  | FINEMAP (LMM) | 0.046 | 0.056 | 0.023 | 0.037 | 0.015 | 0.027 | 0.034 |
|  | FINEMAP (LM) | 0.027 | 0.065 | 0.009 | 0.039 | 0.017 | 0.035 | 0.032 |
|  | FINEMAP-inf (LMM) | 0.024 | 0.028 | 0.011 | 0.017 | 0.006 | 0.018 | 0.017 |
|  | FINEMAP-inf (LM) | 0.023 | 0.070 | 0.006 | 0.043 | 0.014 | 0.017 | 0.029 |
|  | SuSiE (LMM) | 0.109 | 0.051 | 0.040 | 0.038 | 0.024 | 0.032 | 0.049 |
|  | SuSiE (LM) | 0.028 | 0.119 | 0.009 | 0.022 | 0.016 | 0.034 | 0.038 |
|  | SuSiE-inf (LMM) | 0.018 | 0.016 | 0.012 | 0.012 | 0.009 | 0.008 | 0.013 |
|  | SuSiE-inf (LM) | 0.011 | 0.059 | 0.004 | 0.021 | 0.007 | 0.022 | 0.021 |

Note: AUPRC values were calculated by varying PIP thresholds (or PCIP for BFMAP-Forward) from 0 to 1 and computing the area under the resulting precision-recall curves. GWAS summary data were derived from GCTA-MLMA (LMM) or PLINK 2.0 --glm (LM). Results are shown for different sample sizes (5,000 and 10,000 individuals), PVE levels (1% and 4%), and numbers of causal variants (1-3).

**Figure 2.**
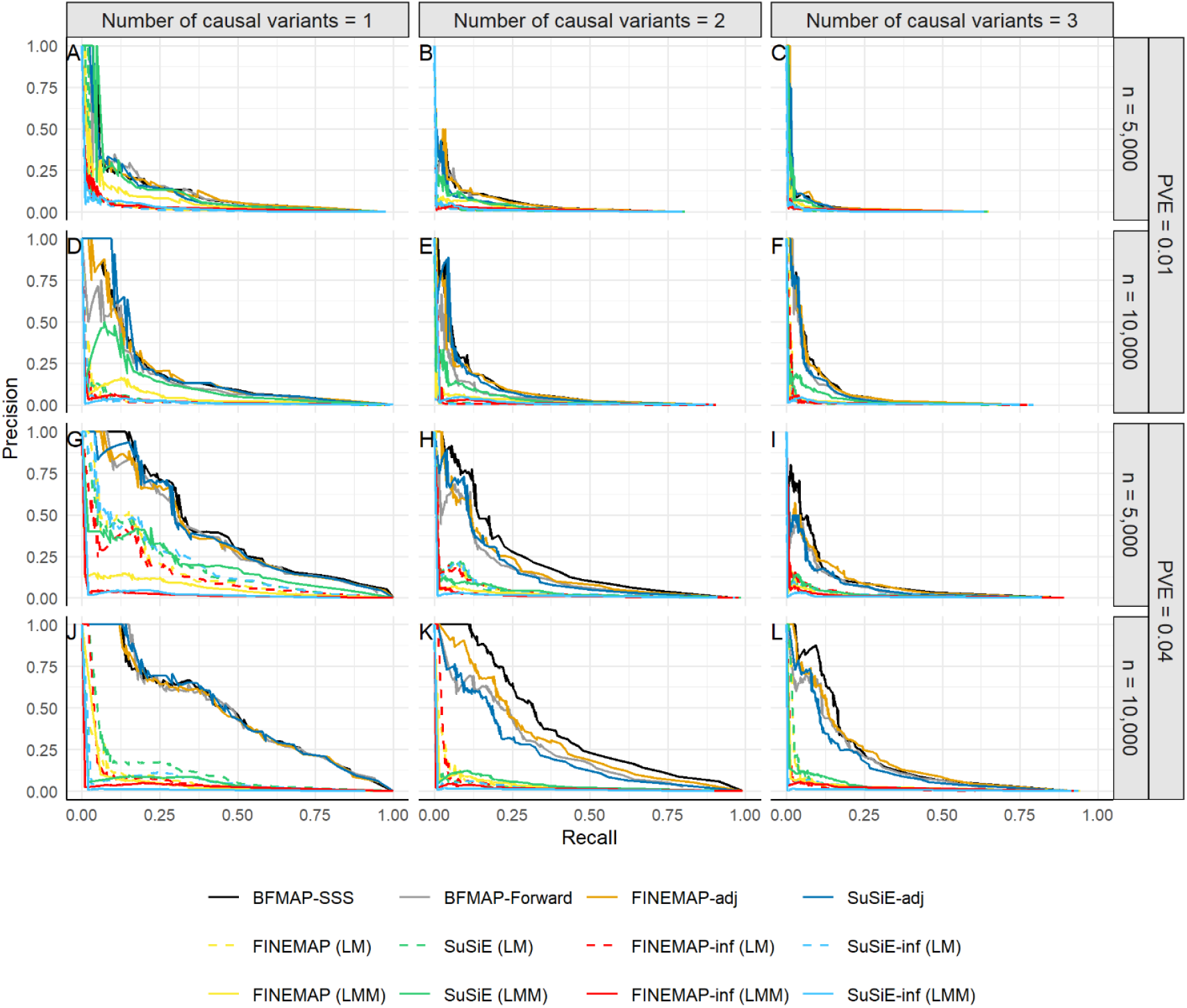
Precision-recall curves for identifying causal variants using PIPs across simulation scenarios. GWAS summary data were derived from GCTA-MLMA (LMM) or PLINK 2.0 --glm (LM). Results are shown for different sample sizes (5,000 and 10,000 individuals), PVE levels (1% and 4%), and numbers of causal variants (1-3), with A-L representing the 12 simulation scenarios.

#### Comparative performance of proposed and existing fine-mapping methods

Overall, GCTA-COJO’s performance in prioritizing causal variants was generally comparable to or slightly poorer than established PIP-based methods across the simulation scenarios (Supplementary Figure S3). Consequently, while its results are available, our primary comparative focus shifts to the PIP/PCIP-based methods.

A consistent pattern emerged when comparing our developed methods (BFMAP-Forward, BFMAP-SSS, FINEMAP-adj, and SuSiE-adj) against the established approaches (FINEMAP, SuSiE, FINEMAP-inf, SuSiE-inf). Our methods designed for related individuals consistently and substantially outperformed these existing tools across all 12 simulation scenarios, as illustrated by the precision-recall curves (Figure 2) and the AUPRC values (Table 1). This performance advantage for our methods was particularly pronounced in scenarios with larger sample sizes or higher PVE. For instance, when AUPRCs were averaged across the six scenarios (3 causal variant counts × 2 PVE levels) with 10,000 individuals, BFMAP-SSS (average AUPRC = 0.234) showed approximately a 3.8-fold higher AUPRC than the best-performing reference method, SuSiE using LMM-derived association statistics (average AUPRC = 0.049) (Table 1). Specifically, FINEMAP-adj and SuSiE-adj, which incorporate relatedness-adjusted LD matrices and LMM-derived association statistics, achieved remarkable improvements over their counterparts (FINEMAP and SuSiE) that use standard LD matrices. When AUPRCs were averaged across the six scenarios (3 causal variant counts × 2 PVE levels) with 10,000 individuals, FINEMAP-adj yielded a 5.4-fold higher AUPRC than standard FINEMAP, and SuSiE-adj a 3.1-fold higher AUPRC than standard SuSiE. These substantial gains were achieved when standard FINEMAP and SuSiE were provided with LMM-derived association statistics, their generally optimal input type under these conditions with 10,000 individuals (Table 1).

Among our LMM-based approaches, the overall differences in fine-mapping performance were relatively small (Table 1, Figure 2). BFMAP-SSS, employing shotgun stochastic search, performed best overall. BFMAP-Forward, which uses a forward selection strategy, showed performance comparable to or slightly less favorable than FINEMAP-adj and SuSiE-adj that implement more exhaustive model space exploration.

#### Fine-mapping challenges in livestock populations and impact of simulation parameters

It is crucial to note the inherent difficulty of fine-mapping in livestock populations, characterized by strong LD and extensive genetic relatedness among individuals. Even with methods explicitly accounting for relatedness and whole-genome infinitesimal effects, performance can be constrained. For instance, in the simulation scenario with 3 causal variants, 10,000 individuals, and 1% PVE, the best-performing reference method (SuSiE, using its optimal input) yielded an AUPRC of only 0.024 (Table 1). While our novel methods offered a substantial improvement (e.g., BFMAP-SSS achieved an AUPRC of 0.077 in the same scenario), these results underscore that fine-mapping in such populations remains a formidable task.

The simulation parameters also highlighted expected trends. Performance for most methods improved with fewer causal variants or larger PVE for the candidate region. A key observation was the differential response to increased sample size: while our relatedness-aware approaches (BFMAP-Forward, BFMAP-SSS, FINEMAP-adj, and SuSiE-adj) uniformly benefited from larger sample sizes, the methods originally designed for unrelated individuals (FINEMAP, SuSiE, and their -inf variants) showed limited improvement or, in some instances, even a decline in performance as sample size increased when applied to these related samples (Table 1). This underscores the critical importance of appropriately modeling genetic relatedness, especially in larger cohorts of related individuals.

### Gene-level PIPs increase detection power in fine-mapping

To address the challenges of fine-mapping at single-variant resolution, particularly in populations with extensive LD like the Duroc pigs in this study, we evaluated the utility of our proposed gene-level posterior inclusion probability (PIP_gene_). The calculation of PIP_gene_ requires posterior model probabilities for variant sets, which are not standard outputs for all fine-mapping tools. Therefore, for this specific evaluation, we computed PIP_gene_ using the outputs from BFMAP-Forward, BFMAP-SSS, and FINEMAP-adj, as these methods provide the necessary model-level information. This selection of methods is sufficient to demonstrate the advantages of gene-level aggregation over variant-level PIPs, an orthogonal comparison to the inter-method performance benchmarks detailed previously.

The application of gene-level fine-mapping resulted in a substantial improvement in the ability to correctly identify causal genes (i.e., our defined gene proxy blocks containing one or more true causal variants) for all three methods evaluated (BFMAP-Forward, BFMAP-SSS, and FINEMAP-adj). This enhancement is evident from the precision-recall curves, which consistently show superior performance for gene-level PIPs compared to variant-level PIPs across all 12 simulation scenarios (Figure 3). The AUPRC values further underscore the benefits of gene-level aggregation (Table 2). The AUPRC for identifying causal genes was markedly higher than that for identifying individual causal variants across all methods and scenarios. Notably, the magnitude of this performance gain was often greatest in the more challenging simulation scenarios. For instance, in the scenario with 3 causal variants, 1% PVE, and 10,000 individuals (a setting where variant-level fine-mapping proved particularly difficult), gene-level aggregation yielded dramatic improvements. At the variant level, BFMAP-Forward, BFMAP-SSS, and FINEMAP-adj achieved AUPRCs of 0.061, 0.077, and 0.076, respectively. In contrast, when evidence was aggregated to the gene level using PIP_gene_, these same methods achieved AUPRCs of 0.305, 0.420, and 0.406, respectively (Table 2). This represents an approximate 4-to 5.5-fold increase in AUPRC, highlighting the substantial power gained by shifting the inferential focus from individual variants to gene units in complex fine-mapping contexts.

**Table 2.** AUPRC for causal gene identification by fine-mapping methods across simulation scenarios.

| Sample size | Method | Number of causal variants |  |  |  |  |  |
| --- | --- | --- | --- | --- | --- | --- | --- |
|  |  | 1 |  | 2 |  | 3 |  |
|  |  | PVE=0.01 | PVE=0.04 | PVE=0.01 | PVE=0.04 | PVE=0.01 | PVE=0.04 |
| 5,000 | BFMAP-SSS | 0.584 | 0.836 | 0.404 | 0.686 | 0.267 | 0.507 |
|  | BFMAP-Forward | 0.535 | 0.780 | 0.277 | 0.548 | 0.149 | 0.371 |
|  | FINEMAP-adj | 0.574 | 0.811 | 0.403 | 0.622 | 0.265 | 0.479 |
| 10,000 | BFMAP-SSS | 0.688 | 0.856 | 0.467 | 0.782 | 0.420 | 0.662 |
|  | BFMAP-Forward | 0.618 | 0.828 | 0.341 | 0.585 | 0.305 | 0.527 |
|  | FINEMAP-adj | 0.693 | 0.852 | 0.455 | 0.721 | 0.406 | 0.635 |

Note: AUPRC values were calculated by varying PIP thresholds (or PCIP for BFMAP-Forward) from 0 to 1 and computing the area under the resulting precision-recall curves. Results show the ability of three methods (BFMAP-Forward, BFMAP-SSS, and FINEMAP-adj) to identify causal genes (defined as gene proxy blocks containing true causal variants) under different sample sizes (5,000 and 10,000 individuals), PVE levels (1% and 4%), and numbers of causal variants (1-3).

**Figure 3.**
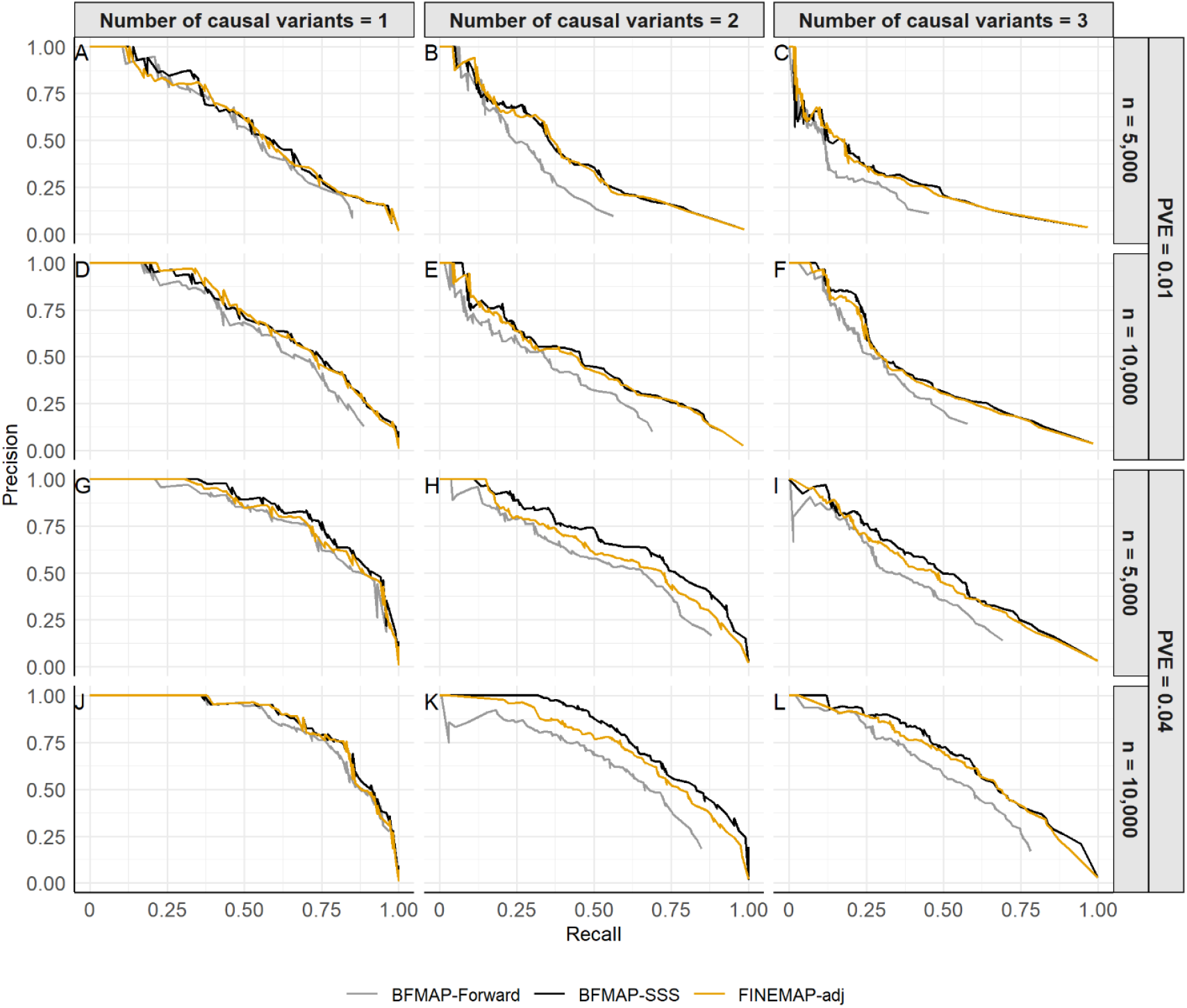
Precision-recall curves for identifying causal genes across simulation scenarios. Results show the ability of three methods (BFMAP-Forward, BFMAP-SSS, and FINEMAP-adj) to identify causal genes (defined as gene proxy blocks containing true causal variants) under different sample sizes (5,000 and 10,000 individuals), PVE levels (1% and 4%), and numbers of causal variants (1-3). A-L represent the 12 simulation scenarios.

### Computational performance

We evaluated the runtime of our LMM-based fine-mapping methods using simulated datasets with 2 causal variants (4% PVE) for 5,000 and 10,000 individuals, focusing on the same 4-Mb candidate region (3,837 and 4,371 pre-filtered SNPs, respectively) as used in our main simulation benchmarks. Analyses were conducted on an Intel Xeon Gold 6230 CPU. Key fine-mapping parameters, such as the prior PVE per causal variant (0.01) and the maximum number of assumed causal variants (5), were kept consistent with the main simulation benchmarks.

Wall-clock times for all methods, representing the analysis of a single simulation replicate, are detailed in Supplementary Table S2. BFMAP-Forward was the fastest approach, completing analyses in approximately 0.3 minutes (5,000 individuals) and 0.5 minutes (10,000 individuals) with 20 CPU threads. FINEMAP-adj required approximately 1.5 minutes (5,000 individuals) and 2.3 minutes (10,000 individuals) using 20 threads; these times include the pre-calculation of the relatedness-adjusted LD matrix. Both BFMAP-Forward and FINEMAP-adj showed limited additional speedup when using 20 threads compared to single-thread execution. BFMAP-SSS, while generally the most accurate in fine-mapping benchmarks, was the most computationally intensive, with wall-clock times of approximately 52 minutes (5,000 individuals) and 170 minutes (10,000 individuals) using 20 threads. SuSiE-adj, which utilizes the susieR package and operates on a single thread, completed its analyses in approximately 1.3 minutes (5,000 individuals) and 2.3 minutes (10,000 individuals), comparable to single-threaded FINEMAP-adj.

### Application to Duroc pig traits

We applied our novel fine-mapping methods to analyze six economically important traits in the Duroc pig population previously described. To identify candidate genomic regions for fine-mapping, we first performed GWAS for each trait using the mixed-model approach implemented in SLEMM-GWA. All six traits produced high-quality Manhattan plots, five of which displayed association peaks with an inclusive significance threshold of *P* < 1 × 10^−5^ (Supplementary Figure S4). Using this threshold, we identified 46 candidate region-trait pairs: 43 for performance traits and 3 for reproduction traits (Supplementary Table S3).

To identify putative causal variants and genes within these candidate regions, we initially employed BFMAP-SSS due to its strong performance in simulations. This fine-mapping identified 30 variants with a PIP > 0.1 and 2 variants with a PIP > 0.5 across all region-trait pairs (Supplementary Table S4). Recognizing the potential to enhance discovery by aggregating variant-level evidence, we then calculated the gene-level PIP (PIP_gene_) for all annotated genes within each candidate region, with gene boundaries defined as detailed in Material and methods. This approach identified 87 gene-trait pairs with PIP_gene_ > 0.1, of which 22 pairs had a PIP_gene_ > 0.5 (Supplementary Table S5). These results underscore the utility of this gene-based strategy for prioritizing candidate genes.

The power of PIP_gene_ was particularly evident in regions where individual variant PIPs were low. For example, a candidate region on chromosome 1 (chr1:51,300,453-55,300,453) associated with back fat thickness (BF) (Figure 4A) contained no variants with a PIP greater than 0.03 (Figure 4B), making variant prioritization challenging. However, the PIP_gene_ analysis revealed that *MRAP2* emerged with a strong signal (PIP_gene_ = 0.669, distinctly higher than other genes in the region; Figure 4C). The *MRAP2* gene encodes melanocortin receptor accessory protein 2, which regulates melanocortin receptor activity and energy homeostasis (31). Previous studies have identified loss-of-function *MRAP2* variants as pathogenic for monogenic hyperphagic obesity in humans (32, 33), and *Mrap2*-knockout mice develop severe early-onset obesity with increased fat mass and adipocyte hypertrophy (33). These findings strongly support *MRAP2* as a biologically plausible candidate for the BF association observed in our study.

**Figure 4.**
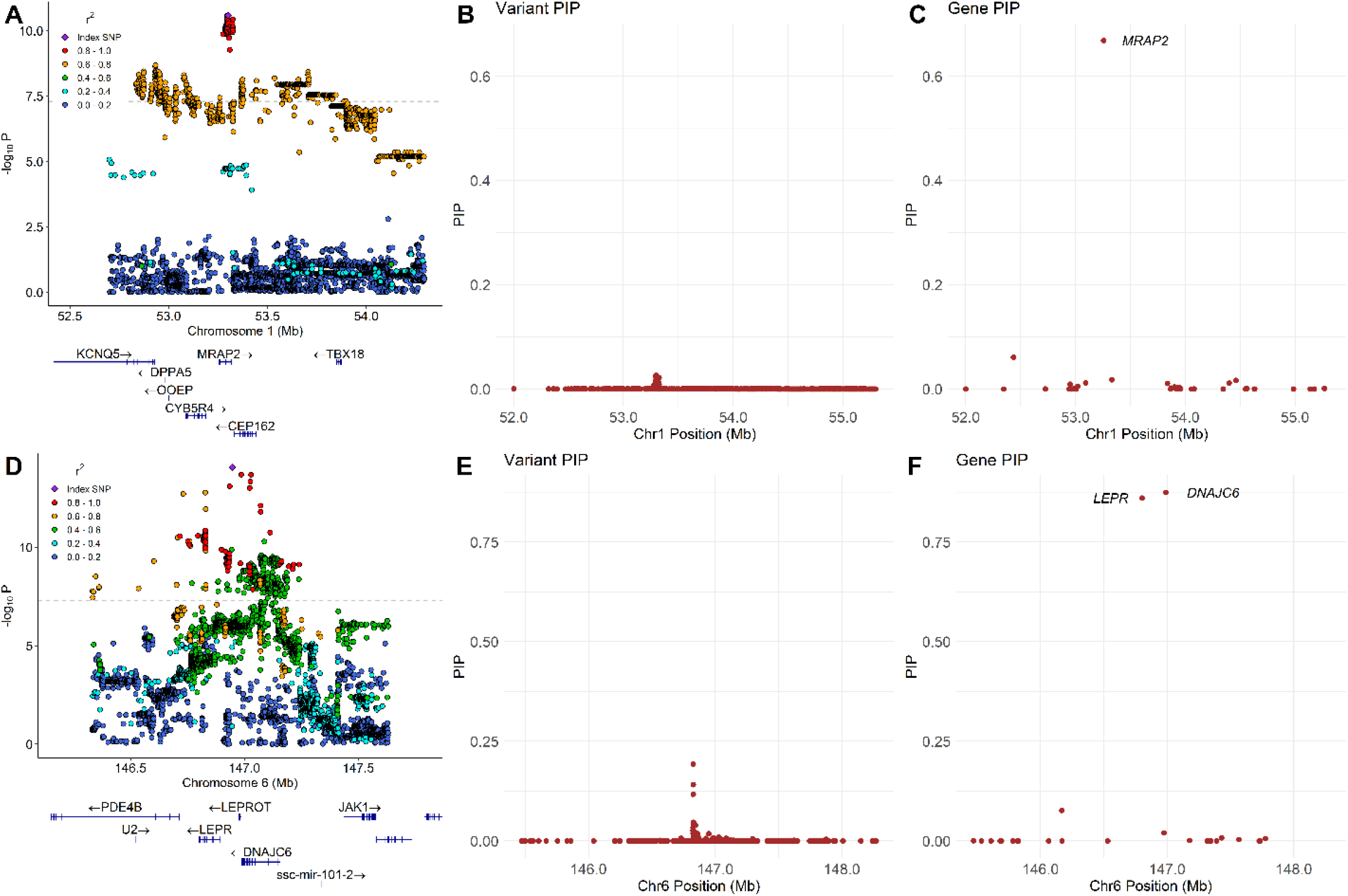
Fine-mapping results for two candidate regions. BFMAP-SSS fine-mapping analysis of two genomic regions associated with back fat thickness: chr1:51,300,453–55,300,453 and chr6:144,948,358–148,948,358. (A) and (D) show locus zoom plots displaying local association signals within each interval. (B) and (E) show the PIPs of individual variants, and (C) and (F) show the PIPs of genes.

A second illustrative example for BF is the candidate region on chromosome 6 (chr6:144,948,358-148,948,358; Figure 4D). In this locus, fine-mapping with BFMAP-SSS identified three prioritized variants (each with PIP > 0.1) that are located within the *LEPR* gene. This gene encodes the leptin receptor, which mediates leptin signaling in energy homeostasis and body weight regulation (34). The *LEPR* gene showed a strong gene-level signal (PIP_gene_ = 0.861), consistent with its well-established role in regulating energy homeostasis and body composition (35–37). Mutations in this gene can cause severe early-onset obesity, hyperphagia, and hypogonadotropic hypogonadism in humans (38, 39). While PIPs of all other variants in the candidate region were < 0.05 (Figure 4E), our PIP_gene_ analysis highlighted an additional candidate gene, *DNAJC6* (PIP_gene_ = 0.875; Figure 4F). This gene encodes auxilin-1, a DNAJ/HSP40 family protein that regulates resting metabolic rate (40) and has been shown to suppress adipogenesis and energy metabolism in adipocytes (41). These findings demonstrate the ability of PIP_gene_ to enhance detection power for relevant genes whose signals might be diffuse at the individual variant level.

Finally, we compared our summary-statistics-based methods (FINEMAP-adj and SuSiE-adj) against individual-level BFMAP-SSS by re-analyzing the two illustrative BF candidate regions. At the variant level, SuSiE-adj PIPs showed high concordance with BFMAP-SSS PIPs in both regions (Supplementary Figures S5A and S5B). For FINEMAP-adj, variant PIPs on the chromosome 1 region were highly concordant with BFMAP-SSS (Supplementary Figure S5A) and stable across iteration lengths (Supplementary Figures S6A and S6C), including the default iteration settings. The chromosome 6 region, however, proved more challenging for FINEMAP-adj: while its concordance with BFMAP-SSS improved with increased iterations from the default settings (plateauing around 10,000 iterations) (Supplementary Figures S6B and S6D), FINEMAP-adj still showed substantial discrepancy from BFMAP-SSS, with a PIP correlation between the two methods of 0.594 even with 50,000 iterations (Supplementary Figure S5B). Despite the variant-level discrepancy for FINEMAP-adj on the chromosome 6 region, PIP_gene_ values derived from the FINEMAP-adj runs with 50,000 iterations were highly correlated with those from BFMAP-SSS in both regions (chr1: 0.999; chr6: 0.945; Supplementary Figure S5C and S5D). Gene-level aggregation yielded more highly consistent inferences between FINEMAP-adj and BFMAP-SSS, even when variant-level concordance is limited (Supplementary Figure S6).

## Discussion

Fine-mapping complex traits in populations with extensive relatedness, such as livestock, presents significant statistical challenges. Standard fine-mapping methodologies, predominantly developed for human genetics studies involving largely unrelated individuals, can falter or yield unreliable results when applied naively to these populations. This study addresses this critical gap by introducing a comprehensive framework. This includes BFMAP (a Bayesian approach for individual-level data, encompassing our previously developed BFMAP-Forward (17) and the novel BFMAP-SSS implementation), innovative summary-statistics adaptations (FINEMAP-adj and SuSiE-adj), and a gene-level aggregation metric (PIP_gene_), all designed to improve fine-mapping under such complex conditions of relatedness.

A key contribution of this work is the clear demonstration, both theoretically and through extensive pig genotype-based simulations, that appropriately modeling relatedness is critical for achieving accurate fine-mapping. Our results unequivocally show that standard fine-mapping tools (GCTA-COJO, FINEMAP, SuSiE, and their -inf variants), even when provided with summary association statistics from mixed models, exhibit substantially poorer performance than our relatedness-aware methods (BFMAP-Forward, BFMAP-SSS, FINEMAP-adj, and SuSiE-adj) designed to correctly utilize relatedness information (Figure 2, Table 1). The performance gap, reflected in markedly poorer precision-recall profiles and lower AUPRC values for existing methods (GCTA-COJO, FINEMAP, SuSiE, and their -inf variants) (Figure 2, Table 1), underscores the potential for a high proportion of false positives in previous fine-mapping studies in livestock that applied human-centric fine-mapping methods not designed for samples of related individuals (e.g., 42–46). It is important to note that simply §using LMM-derived summary association statistics was insufficient to overcome the fundamental limitations of these methods when applied to samples of related individuals. This highlights the critical need for approaches that explicitly integrate relatedness, primarily through a relatedness-adjusted LD matrix, as implemented in our -adj methods, or via direct LMM modeling as in BFMAP.

Among our proposed relatedness-aware methods, BFMAP-SSS generally emerged as the most accurate, likely due to its sophisticated shotgun stochastic search with simulated annealing allowing for a more thorough exploration of the model space. FINEMAP-adj and SuSiE-adj also performed well, trailing BFMAP-SSS and slightly outperforming BFMAP-Forward (Table 1, Figure 2). This performance hierarchy aligns with theoretical expectations: the forward selection strategy of BFMAP-Forward is inherently less exhaustive than the search algorithms in BFMAP-SSS or the variable selection approaches underpinning SuSiE (and thus SuSiE-adj). The slight performance edge of BFMAP-SSS over the -adj methods may stem from the GRAMMAR-Gamma-like approximation employed to allow the FINEMAP and SuSiE programs to be directly applied using appropriately transformed summary statistics (LMM-derived z-scores and effective sample size *ñ*) and a relatedness-adjusted LD matrix 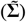. While developing new programs for FINEMAP-adj and SuSiE-adj from scratch to circumvent this approximation is feasible, the current -adj approaches offer substantial practical advantages by leveraging well-established, existing software, especially given their performance close to BFMAP-SSS.

The choice among our developed methods in practice will depend on a balance of factors. BFMAP-SSS offers the highest accuracy but is the most computationally intensive. For studies where computational resources are a constraint, FINEMAP-adj, SuSiE-adj, and BFMAP-Forward provide much faster alternatives with only a modest trade-off in accuracy compared to BFMAP-SSS (Figure 2, Supplementary Table S2). FINEMAP-adj, BFMAP-SSS, and BFMAP-Forward also readily provide the posterior model probabilities necessary for calculating PIP_gene_, a feature not directly available from SuSiE-adj (which builds on SuSiE) without further methodological development. Furthermore, users should be aware that FINEMAP-adj (which builds on FINEMAP) can sometimes require extended iterations for convergence in complex regions, whereas SuSiE-adj, benefiting from SuSiE’s efficient variable selection, is generally robust in convergence. We therefore recommend BFMAP-SSS when maximal accuracy is paramount and resources permit; FINEMAP-adj when gene-level insights are desired with good speed; SuSiE-adj for fast and robust variant-level fine-mapping without immediate need for PIP_gene_; and BFMAP-Forward as a rapid option for individual-level data.

A significant challenge in livestock genomics is the extensive and complex LD structure, usually making fine-mapping at single-variant resolution exceedingly difficult. Our proposed PIP_gene_ metric addresses this by aggregating variant-level evidence, thereby enhancing the power to detect trait-associated genes, albeit with a trade-off in localization precision from variant to gene. As demonstrated in our simulations, the improvement in detection power (AUPRC) using PIP_gene_ was substantial, particularly in challenging scenarios with low PVE and multiple causal variants, where gene-level AUPRCs were up to 5.5-fold higher than variant-level AUPRCs (Table 2, Figure 3). This utility was further underscored in our analysis of Duroc pig traits, where PIP_gene_ successfully highlighted biologically relevant genes such as *MRAP2* and *DNAJC6* for back fat thickness, even when underlying variant PIPs were diffuse or individually weak (Figure 4). Furthermore, this approach led to the identification of other promising candidate genes, such as *DYRK4* as a novel candidate for back fat thickness, whose high PIP_gene_ values warrant further investigation (47). These findings collectively highlight the value of our fine-mapping framework for enhancing candidate gene discovery in livestock.

Despite the advancements presented, this study has limitations. A primary limitation is our simulation design. While based on real pig genotypes, evaluations focused on a single 4-Mb candidate region where we simulated only 1-3 variants as having larger causal effects. This, combined with an infinitesimal background modeled by 10,000 genome-wide chip SNPs, represents one specific model of genetic architecture; real traits exhibit greater heterogeneity in the number, effect size distribution, and genomic location of causal variants. Furthermore, the GRM used to model this infinitesimal background was constructed directly from these 10,000 known infinitesimal-effect SNPs and was then supplied as input to GRM-dependent fine-mapping methods. This perfect concordance between the SNPs defining the simulated infinitesimal background and those in the analytical GRM is an idealization not reflective of real-world scenarios, where GRMs are estimated from available marker data (often LD-pruned) and may not fully capture true genetic covariance or perfectly align with the underlying polygenic architecture. While our main simulations effectively highlighted performance differences using this consistent GRM, we further investigated the impact of using a more “practical” GRM, constructed from 10,000 randomly selected quality-controlled genome-wide variants (from the ∼11.7 million). Re-analyzing all simulation scenarios with FINEMAP-adj and SuSiE-adj using this practical GRM (for both LMM-GWAS summary statistic generation and as input to the -adj methods) yielded fine-mapping performance highly concordant with that obtained using the “simulation-derived” GRM (Supplementary Figure S7, Supplementary Table S6). This result suggests that while GRM misspecification remains a theoretical concern, our -adj approaches demonstrate robustness to this aspect of GRM construction, at least with reasonably dense marker-based GRMs. While our study effectively demonstrates the advantages of relatedness-aware methods under the tested architectures, further research exploring a wider array of simulated genetic complexities will be valuable for comprehensively understanding their performance boundaries across all potential real-world scenarios. Secondly, the current implementations, particularly those involving matrix operations on the GRM or relatedness-adjusted LD, may face scalability challenges with datasets involving hundreds of thousands of individuals, a scale becoming increasingly common. Further algorithmic refinements will be needed to cater to such massive datasets. Thirdly, our current PIP_gene_ formulation, which sums posterior model probabilities for models including any variant within a gene region, can be systematically influenced by gene length and variant density. Under the standard fine-mapping prior that assigns equal *a priori* causal probability to each variant, longer genes with more variants inherently have greater opportunities to accumulate signal, leading to higher PIP_gene_ values that may not solely reflect biological enrichment for causality. Future refinements could explore incorporating more sophisticated priors that account for gene size or variant density to further enhance the specificity of gene-level inference. Fourthly, our real data analysis relied on imputed sequence data for SNPs and indels; other structural variations were not considered, and differential imputation quality across variants could introduce unmodeled uncertainty.

Future research could focus on several exciting avenues. Incorporating functional annotation data into the prior model probabilities for variants or genes (17, 48–50) holds considerable promise for further enhancing the statistical power and accuracy of our methods. Additionally, while this study focused on fine-mapping GWAS signals for complex traits, the principles of accounting for relatedness (particularly through adjusted LD matrices and mixed models) are broadly applicable. Investigating the performance and adaptation of our framework for other types of genetic analyses in samples of related individuals, such as fine-mapping expression quantitative trait loci (eQTLs) (51–54), represents a valuable direction.

In conclusion, this study provides a robust suite of fine-mapping tools (BFMAP-Forward, BFMAP-SSS, FINEMAP-adj, and SuSiE-adj) and a novel gene-level aggregation strategy (PIP_gene_) specifically designed for and validated in populations with complex relatedness, such as livestock. By properly accounting for genetic relationships and whole-genome infinitesimal effects, these methods offer substantial improvements in accuracy and power over existing approaches, paving the way for more reliable identification of causal variants and genes underlying complex traits in these important populations.

## Acknowledgments

We appreciate Smithfield Premium Genetics (Rose Hill, NC, USA) for providing access to the data used in this study.

## Author contributions

J.J. conceived and designed the study and developed the methodologies. J.Wang carried out the simulation studies and pig trait analyses. F.T., Y.H., and C.M. contributed to data preprocessing. J.Wang, J.J., and J.Wei performed data visualization. J.J. and J.Wang wrote the manuscript. All authors reviewed and approved the final manuscript.

## Conflict of interest

Y.H. was employed by Smithfield Premium Genetics. The remaining authors declare that the research was conducted in the absence of any commercial or financial relationships that could be construed as a potential conflict of interest.

## Funding

This work is supported by the Agriculture and Food Research Initiative (AFRI) Foundational and Applied Science Program, project award no. 2023-67015-39260, and the Research Capacity Fund (HATCH), project award no. 7008128, from the U.S. Department of Agriculture’s National Institute of Food and Agriculture.

## Data availability

BFMAP can be downloaded at https://github.com/jiang18/bfmap. Tutorials for FINEMAP-adj and SuSiE-adj are available at https://github.com/JJWang259/FineMapping-RelatedIndividuals. The Duroc pig data used in this study are property of Smithfield Premium Genetics (Rose Hill, NC, USA). Restrictions apply to the availability of these data, which were used under license for the current study and are not publicly available. Requests to access these datasets may be granted under a negotiated research agreement and should be directed to Kent Gray, General Manager.

## Notes

### Summary of Updates

This version represents a substantial revision of the manuscript with significant updates throughout. The core adapted summary-statistics-based fine-mapping methodology remains unchanged, but the paper has been extensively rewritten with expanded analyses, additional simulation studies, revised figures, and updated supplementary materials.

