## Supplementary Data for "Fine-mapping methods for complex traits: essential adaptations for samples of related individuals"

### Supplementary Figures

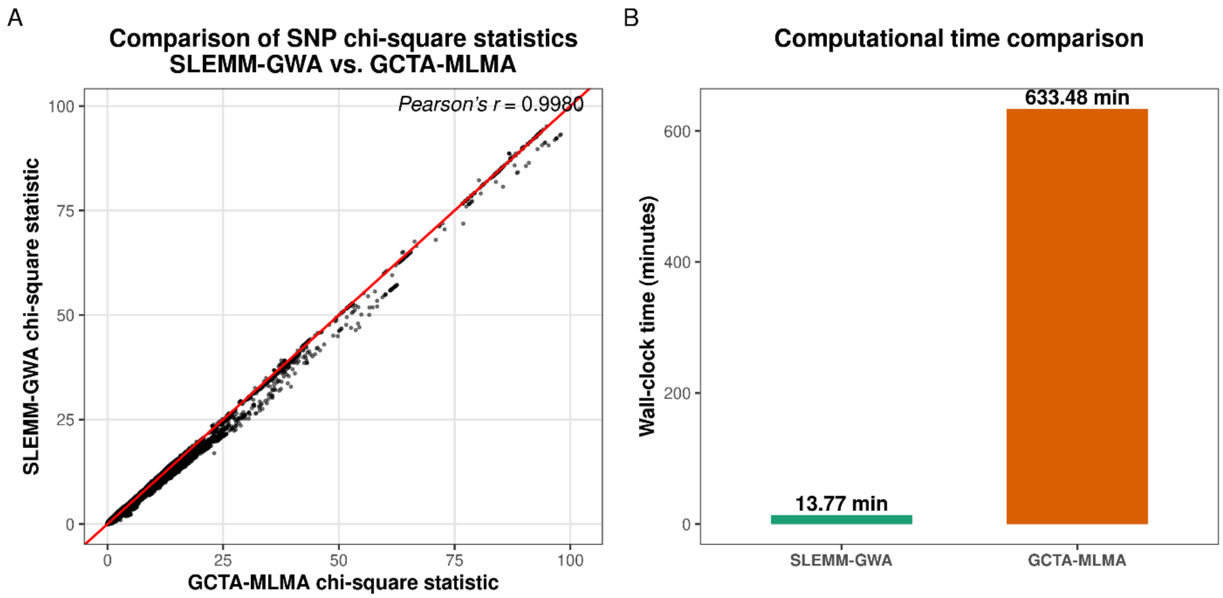


Figure S1. Comparison of GCTA-MLMA and SLEMM-GWA for GWAS of off-test back fat thickness in 26,731 Duroc pigs. Both methods used identical GRMs constructed from 10,000 randomly selected chip SNPs and analyzed 1,000,000 randomly selected imputed variants. (A) Chi-square association test statistic comparison. (B) Computational time comparison. Analyses were conducted on a Linux server with 2×Intel Xeon Gold 6258R CPUs and 512 GB memory using 20 threads.

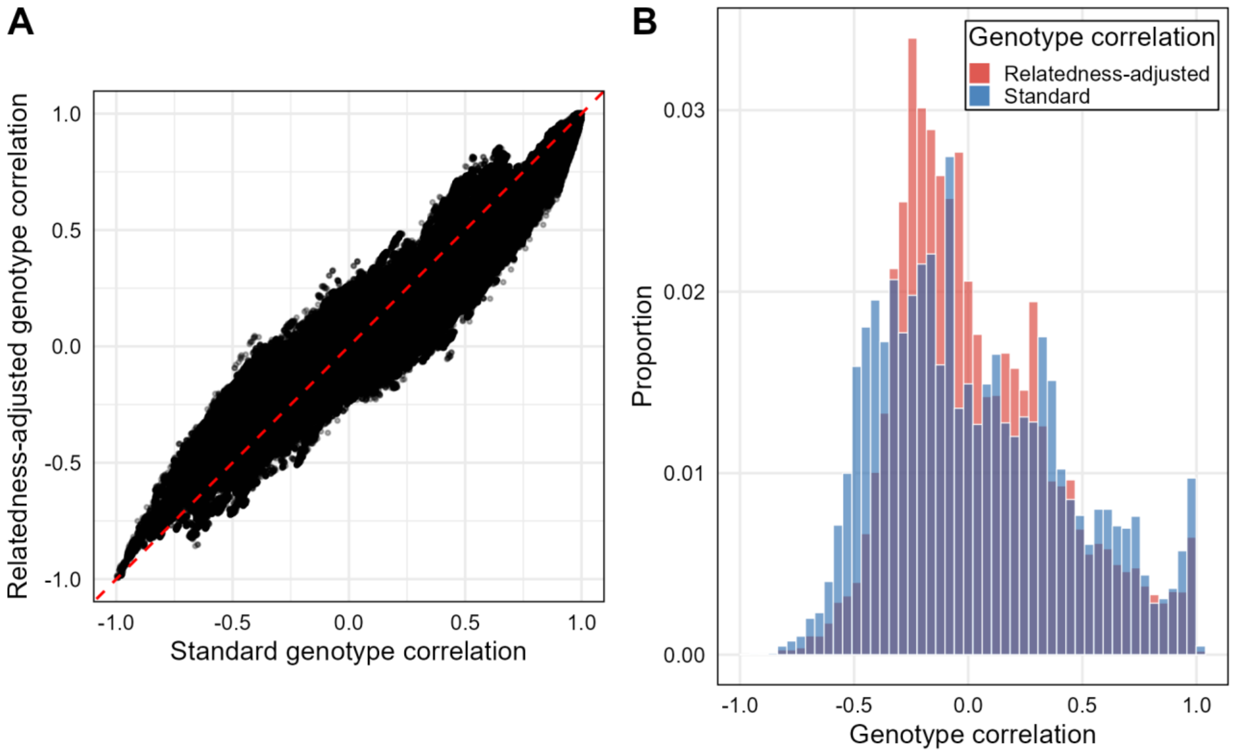


Figure S2. Comparison of standard and relatedness-adjusted genotype correlation matrices. (A) Scatter plot comparing pairwise genotype correlation coefficients between the standard matrix ($\Sigma$) and relatedness-adjusted matrix ($\tilde{\Sigma}$) for all SNP pairs. The red dashed line represents y = x. (B) Density distributions of genotype correlation coefficients for both matrix types. Both matrices were computed using 5,000 randomly selected individuals and 2,456 pre-filtered variants on the chr1 region from one simulation replicate. The standard matrix ($\Sigma$) represents conventional SNP correlations, while the adjusted matrix ($\tilde{\Sigma}$) accounts for genetic relatedness among individuals as described in model (9) with an input heritability parameter of 0.494.

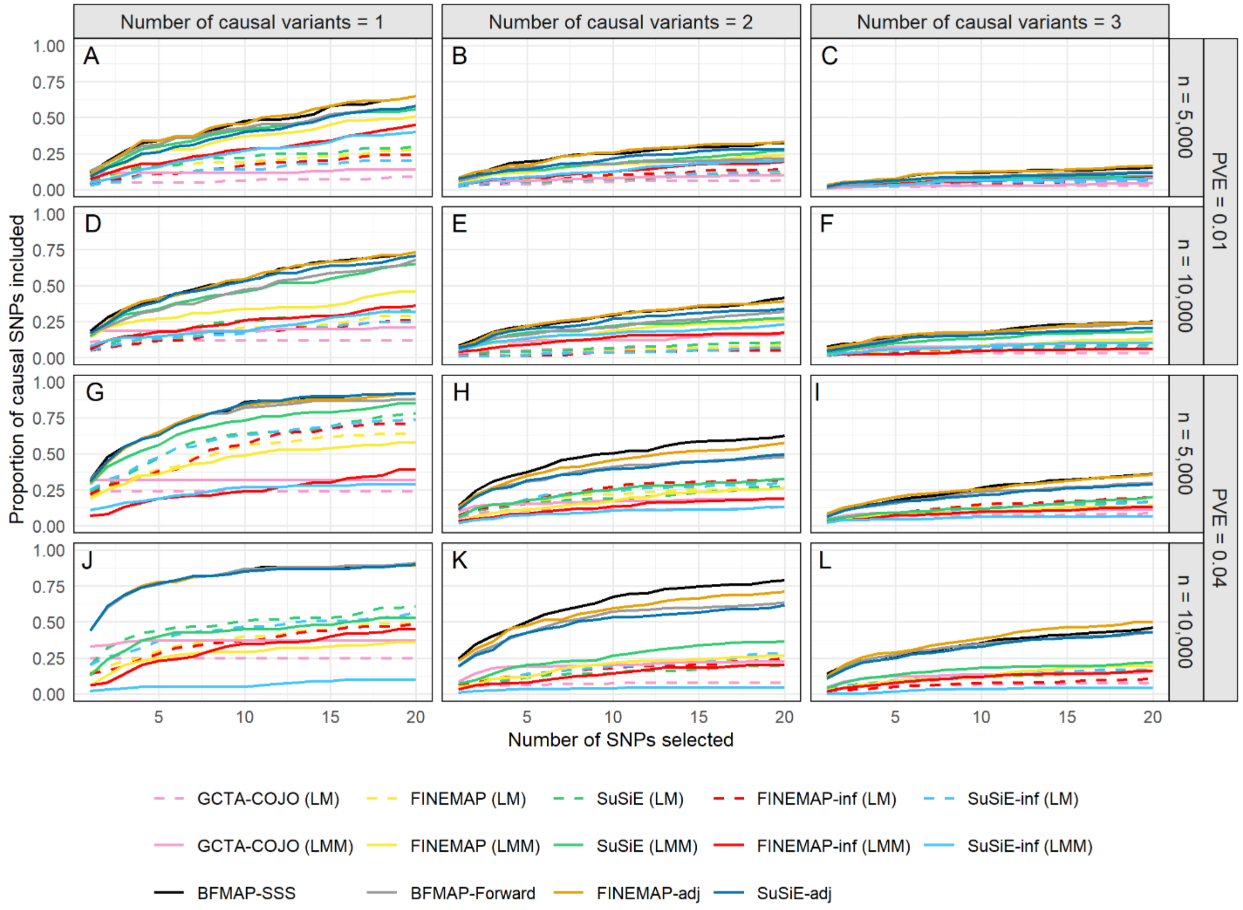


Figure S3. Comparison of fine-mapping methods for identifying causal variants across simulation scenarios. The y-axis shows the proportion of true causal variants correctly identified when selecting the top *k*-ranked variants (x-axis, *k* = 1 to 20) across 12 simulation scenarios. Results are shown for different sample sizes (5,000 and 10,000 individuals), PVE levels (1% and 4%), and numbers of causal variants (1-3). Panels A-L represent the 12 simulation scenarios

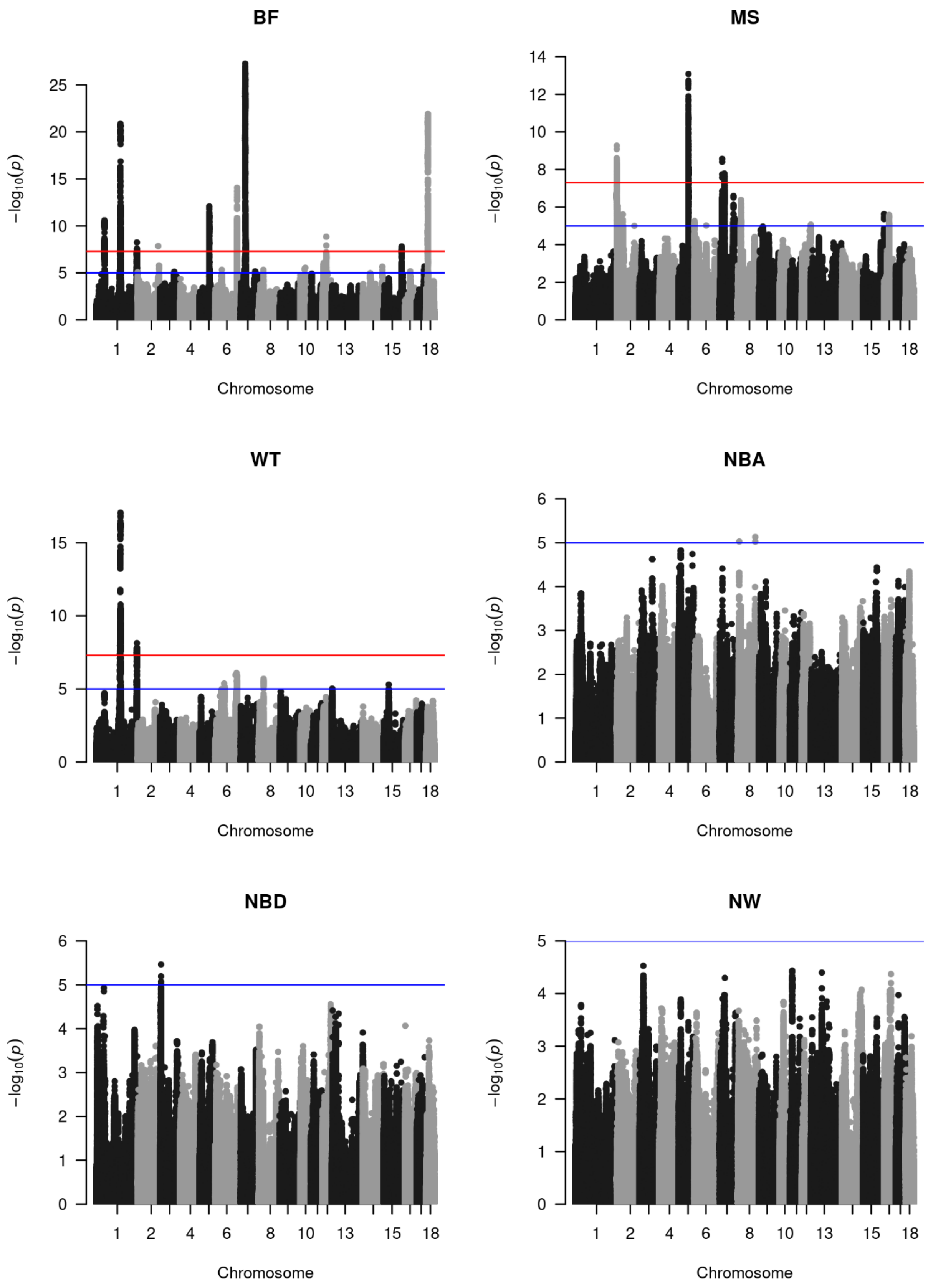
BF
MS
WT
NBA
NBD
NW
$-\log_{10}(p)$
Chromosome
1
2
4
6
8
10
13
15
18

Figure S4. Manhattan plots for GWAS of six quantitative traits in Duroc pigs. Manhattan plots showing GWAS results for three growth traits (off-test body weight [WT], back fat thickness [BF], and loin muscle depth [MS]) analyzed in ~27,000 pigs and three reproduction traits (number of piglets born alive [NBA], born dead [NBD], and weaned [NW]) analyzed in 3,290 sows. Association analyses were performed using SLEMM-GWA with ~11.7 million quality-controlled autosomal variants. GRMs were constructed from 10,000 randomly selected chip SNPs. Horizontal lines indicate genome-wide significance thresholds ($P < 5 \times 10^{-8}$, red) and suggestive significance thresholds ($P < 1 \times 10^{-5}$, blue).

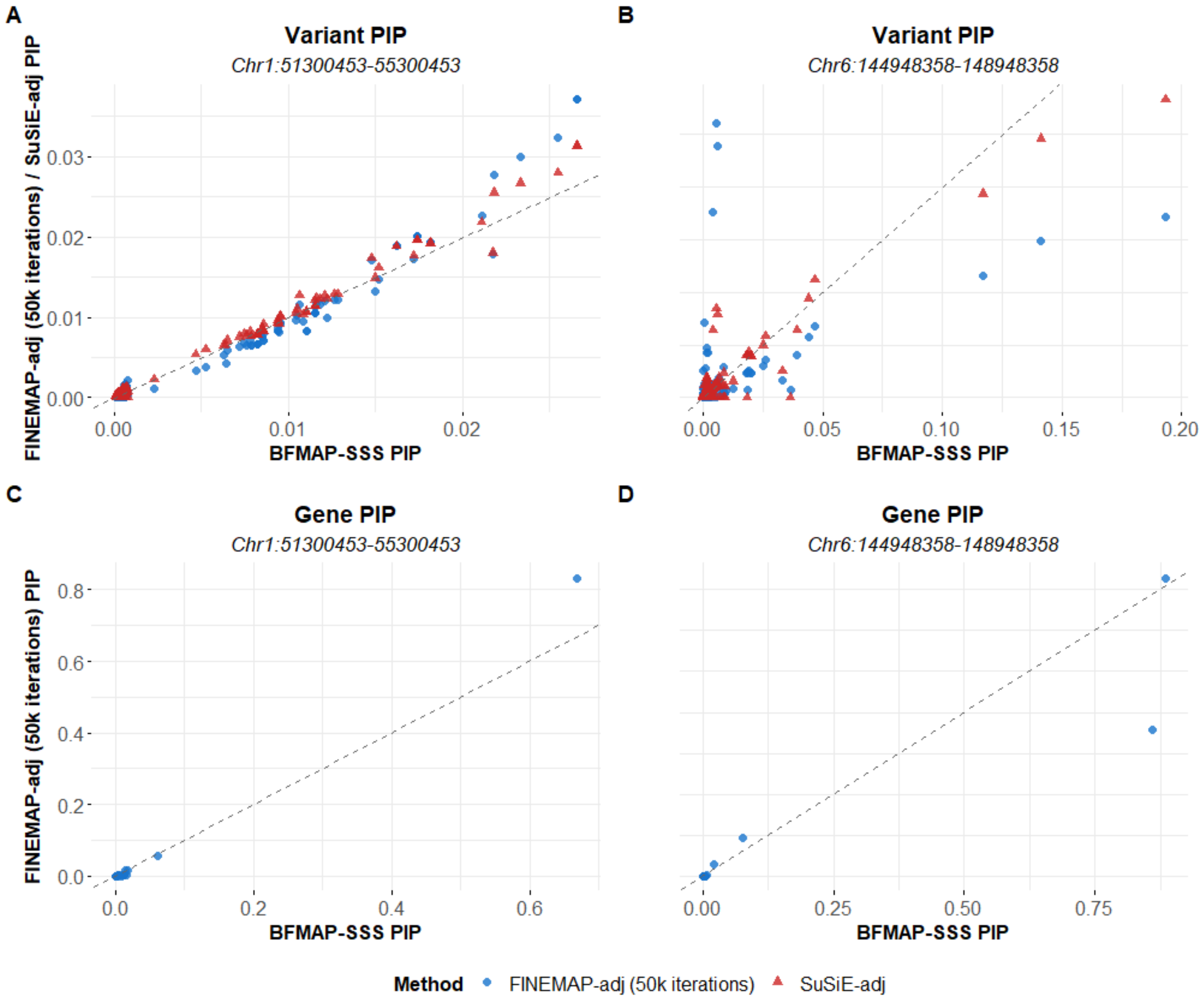


Figure S5. Comparison of PIPs between BFMAP-SSS and adapted summary-statistics-based fine-mapping methods across two genomic regions. Scatter plots comparing variant-level (A, B) and gene-level (C, D) PIPs between BFMAP-SSS and adapted summary-statistics methods for back fat thickness. (A) and (C) show results from chr1:51,300,453-55,300,453 including 4,350 variants. (B) and (D) show results from chr6:144,948,358-148,948,358 including 5,195 variants. The dashed line represents y = x.

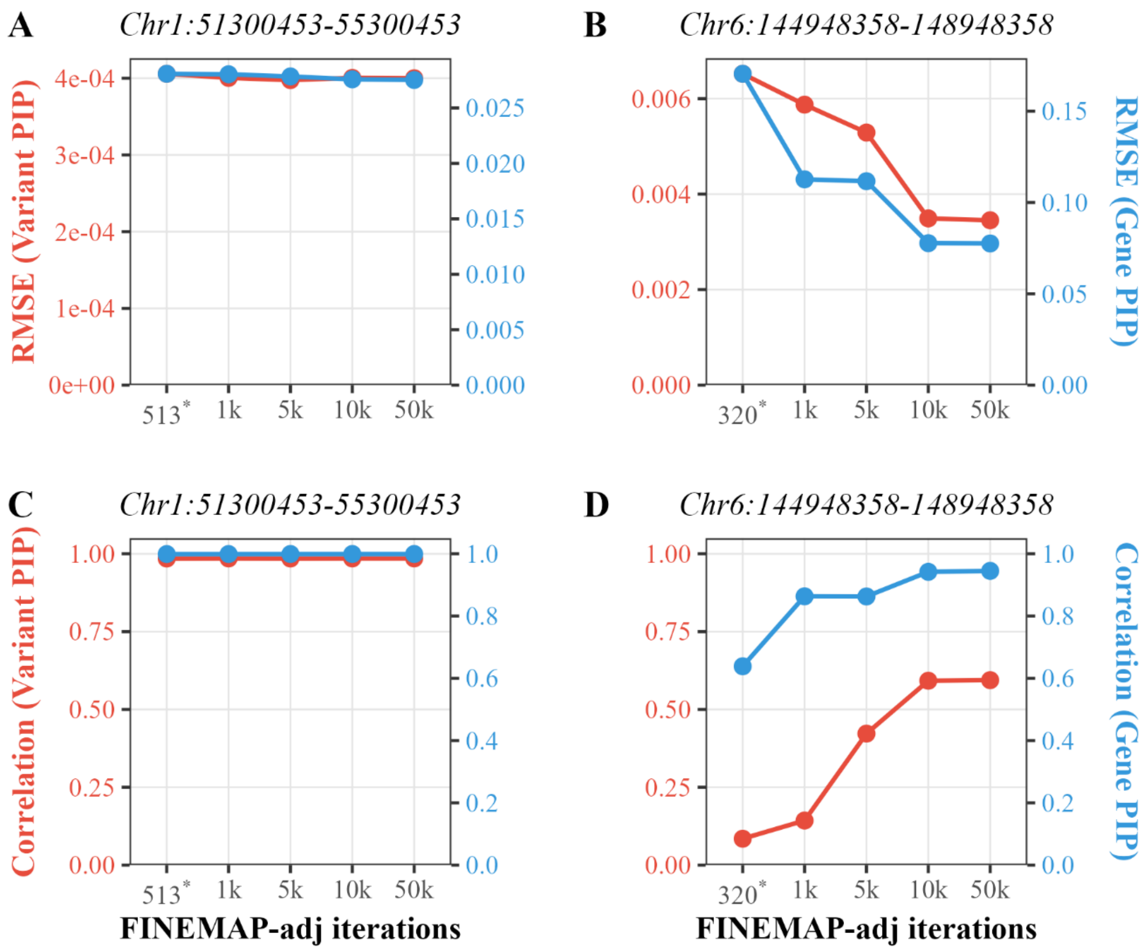


Figure S6 Impact of FINEMAP-adj iteration lengths on PIPs across two genomic regions. Root mean square error (RMSE) (A, B) and correlation (C, D) of PIPs calculated between BFMAP-SSS and FINEMAP-adj at variant level (red) and gene level (blue) across different iteration settings. (A) and (C) show results from chr1:51,300,453-55,300,453, and (B) and (D) from chr6:144,948,358-148,948,358. The x-axis represents FINEMAP-adj iteration numbers. Asterisks (*) indicate actual iteration numbers when applying default convergence settings.

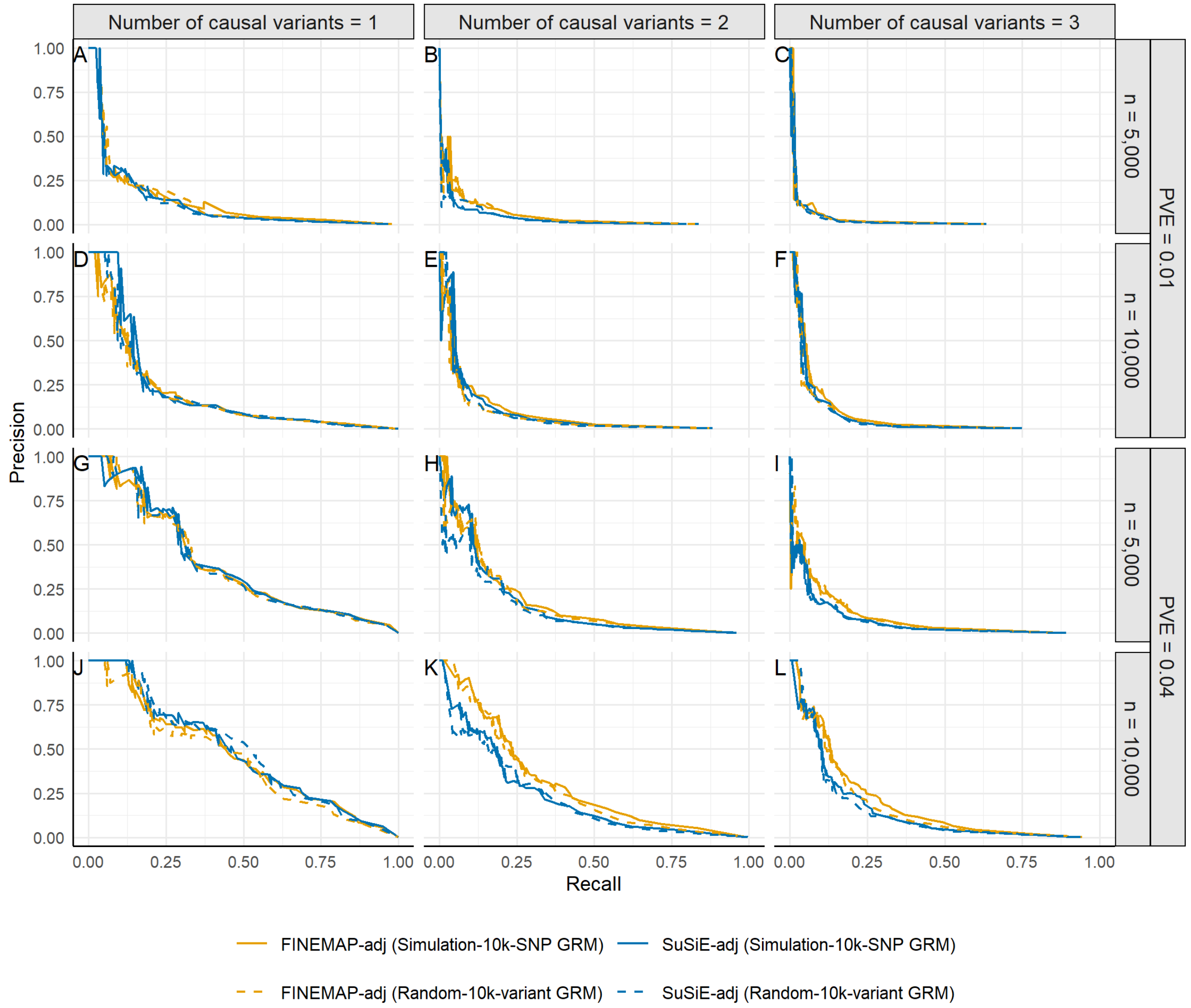


Figure S7. Impact of GRM construction on fine-mapping accuracy. Precision-recall curves comparing FINEMAP-adj and SuSiE-adj when using different GRMs for fine-mapping. Solid lines represent analyses using the same 10,000 chip SNPs used to construct the GRM during the trait simulation (Simulation-10k-SNP GRM), while dashed lines represent analyses using a different set of 10,000 randomly selected variants from the imputed sequence data (Random-10k-variant GRM). Results are shown under different sample sizes (5,000 and 10,000 individuals), PVE levels (1% and 4%), and numbers of causal variants (1-3). Panels A-L correspond to the 12 simulation scenarios.

Table S1. Number of replicates (out of 100) with significant associations ($P < 1\times10^{-5}$) identified by GCTA-MLMA across simulation scenarios.

| Sample size | Number of causal variants | | | | | |
|---|---|---|---|---|---|---|
| | 1 | | 2 | | 3 | |
| | PVE=0.01 | PVE=0.04 | PVE=0.01 | PVE=0.04 | PVE=0.01 | PVE=0.04 |
| 5,000 | 86 | 100 | 74 | 100 | 74 | 98 |
| 10,000 | 97 | 100 | 90 | 100 | 88 | 100 |

Table S2. Computational time (minutes) of LMM-based fine-mapping methods.

| Method | No. of threads | Sample size | |
|---|---|---|---|
| | | 5,000 | 10,000 |
| BFMAP-Forward | 20 | 0.26 | 0.51 |
| FINEMAP-adj | 20 | 1.53 | 2.28 |
| BFMAP-SSS | 20 | 51.83 | 170.02 |
| SuSiE-adj | 1 | 1.30 | 2.30 |
| BFMAP-Forward | 1 | 0.36 | 0.62 |
| FINEMAP-adj | 1 | 1.57 | 2.36 |

Note: Computational time represents wall-clock time for simulation scenario with 2 causal variants explaining 4% of phenotypic variance in a 4-Mb region. Analyses conducted on Intel Xeon Gold 6230 CPU using 20 threads, except SuSiE-adj which is single-threaded. Times for FINEMAP-adj and SuSiE-adj include relatedness-adjusted LD matrix calculation. All methods used prior PVE = 0.01 and maximum 5 causal variants.

Table S3. Candidate regions identified by GWAS for six traits in Duroc pigs.

| Trait | Chromosome | Start | End |
|---|---|---|---|
| BF | 1 | 51300453 | 55300453 |
| BF | 1 | 158383292 | 162383292 |
| BF | 1 | 268769631 | 272769631 |
| BF | 2 | 0 | 3207568 |
| BF | 2 | 137588534 | 141588534 |
| BF | 2 | 142769791 | 146769791 |
| BF | 3 | 92840651 | 96840651 |
| BF | 5 | 63865952 | 67865954 |
| BF | 6 | 42957184 | 46957184 |
| BF | 6 | 144948358 | 148948358 |
| BF | 7 | 22016716 | 26016716 |
| BF | 7 | 28377965 | 32377965 |
| BF | 8 | 28484981 | 32484981 |
| BF | 10 | 26650996 | 30650996 |
| BF | 12 | 5410845 | 9410845 |
| BF | 12 | 22742325 | 26742325 |
| BF | 12 | 28826279 | 32826279 |
| BF | 14 | 128640053 | 132640055 |
| BF | 15 | 116942310 | 120942310 |
| BF | 16 | 33783195 | 37783195 |
| BF | 17 | 48845265 | 52845265 |
| BF | 18 | 8677728 | 12692103 |
| MS | 2 | 0 | 3053429 |
| MS | 2 | 5390887 | 9390887 |
| MS | 2 | 39641510 | 43641510 |
| MS | 5 | 63985520 | 67985520 |
| MS | 6 | 170539 | 4170539 |
| MS | 7 | 13893002 | 17893002 |
| MS | 7 | 28433109 | 32433109 |
| MS | 7 | 90119614 | 94119614 |
| MS | 7 | 95580473 | 99580473 |
| MS | 8 | 18060535 | 22060535 |
| MS | 12 | 59035072 | 63035072 |
| MS | 15 | 137885390 | 141885390 |
| MS | 16 | 31186489 | 35186489 |
| WT | 1 | 157355359 | 162276786 |
| WT | 1 | 270587372 | 270879521 |
| WT | 6 | 60685311 | 64123636 |
| WT | 6 | 137176904 | 137176904 |
| WT | 6 | 143326392 | 143558798 |
| WT | 8 | 31994621 | 32000675 |
| WT | 13 | 3599635 | 3627724 |
| WT | 15 | 32086828 | 32086828 |
| NBA | 8 | 8835736 | 8835736 |

| | | | |
|---|---|---|---|
| NBA | 8 | 116104792 | 116104803 |
| NBD | 3 | 6133790 | 6146253 |

Note: Genomic regions showing significant associations ($P < 1 \times 10^{-5}$) for growth traits (off-test body weight [WT], back fat thickness [BF], and loin muscle depth [MS]) and reproduction traits (number of piglets born alive [NBA], born dead [NBD], and weaned [NW]). Region boundaries were defined by extending 2 Mb upstream and downstream from the position of the minimum *P*-value variant within each associated locus.

Table S4. Candidate variants with PIP > 0.1 identified by BFMAP-SSS.

| Chromosome | Position | PIP | Trait | Candidate region | *P*-value |
|---|---|---|---|---|---|
| 1 | 270945271 | 0.107 | BF | 1:268769631-272769631 | 1.96E-04 |
| 2 | 139588534 | 0.897 | BF | 2:137588534-141588534 | 1.40E-08 |
| 2 | 144907540 | 0.117 | BF | 2:142769791-146769791 | 2.96E-06 |
| 2 | 143121619 | 0.110 | BF | 2:142769791-146769791 | 2.52E-05 |
| 2 | 143121629 | 0.110 | BF | 2:142769791-146769791 | 2.52E-05 |
| 6 | 146826716 | 0.193 | BF | 6:144948358-148948358 | 1.15E-06 |
| 6 | 146826648 | 0.141 | BF | 6:144948358-148948358 | 1.74E-06 |
| 6 | 146826502 | 0.117 | BF | 6:144948358-148948358 | 1.99E-06 |
| 7 | 30377965 | 0.103 | BF | 7:28377965-32377965 | 5.43E-28 |
| 10 | 28650996 | 0.252 | BF | 10:26650996-30650996 | 3.46E-06 |
| 10 | 28649770 | 0.188 | BF | 10:26650996-30650996 | 4.60E-06 |
| 10 | 26814123 | 0.108 | BF | 10:26650996-30650996 | 1.77E-04 |
| 12 | 24901177 | 0.393 | BF | 12:22742325-26742325 | 1.22E-08 |
| 12 | 24742325 | 0.271 | BF | 12:22742325-26742325 | 1.43E-09 |
| 12 | 24028127 | 0.106 | BF | 12:22742325-26742325 | 3.15E-04 |
| 12 | 30826279 | 0.432 | BF | 12:28826279-32826279 | 8.41E-06 |
| 12 | 30826276 | 0.418 | BF | 12:28826279-32826279 | 8.48E-06 |
| 12 | 29199922 | 0.216 | BF | 12:28826279-32826279 | 2.03E-05 |
| 5 | 65985520 | 0.159 | MS | 5:63985520-67985520 | 8.23E-14 |
| 5 | 65752811 | 0.109 | MS | 5:63985520-67985520 | 1.91E-13 |
| 7 | 15893002 | 0.169 | MS | 7:13893002-17893002 | 2.69E-09 |
| 7 | 15908484 | 0.119 | MS | 7:13893002-17893002 | 3.94E-09 |
| 7 | 92119614 | 0.119 | MS | 7:90119614-94119614 | 2.52E-07 |
| 6 | 137176904 | 0.720 | WT | 6:135176904-139176904 | 1.12E-06 |
| 13 | 1816334 | 0.148 | WT | 13:1627724-5627724 | 1.72E-04 |
| 13 | 1815864 | 0.106 | WT | 13:1627724-5627724 | 3.03E-04 |
| 8 | 8835736 | 0.153 | NBA | 8:6835736-10835736 | 9.42E-06 |
| 8 | 8021935 | 0.103 | NBA | 8:6835736-10835736 | 7.64E-04 |
| 8 | 116104792 | 0.298 | NBA | 8:114104792-118104792 | 7.41E-06 |
| 8 | 116104803 | 0.241 | NBA | 8:114104792-118104792 | 9.47E-06 |

Note: *P*-values correspond to single-SNP association statistics from GWAS using SLEMM-GWA. Trait abbreviations: WT, off-test body weight; BF, back fat thickness; MS, loin muscle depth; NBA, number of piglets born alive; NBD, born dead; NW, weaned.

Table S5. Candidate genes with PIP > 0.5 identified by gene-level fine-mapping with BFMAP-SSS.

| Gene | Trait | Chr | Start | End | Gene-PIP | Minimal *P*-value |
|---|---|---|---|---|---|---|
| *MRAP2* | BF | 1 | 53256006 | 53318107 | 0.669 | 2.62E-11 |
| *ABL1* | BF | 1 | 270761668 | 270906708 | 0.637 | 5.75E-09 |
| *NR3C1* | BF | 2 | 144822939 | 144956451 | 0.643 | 2.96E-06 |
| *PRKCE* | BF | 3 | 94349889 | 94866306 | 0.943 | 7.59E-06 |
| *DYRK4* | BF | 5 | 65812243 | 65870092 | 0.676 | 8.69E-13 |
| *DNAJC6* | BF | 6 | 146989351 | 147155131 | 0.884 | 2.10E-14 |
| *LEPR* | BF | 6 | 146801954 | 146895995 | 0.861 | 1.69E-13 |
| *KLHL5* | BF | 8 | 30361338 | 30440750 | 0.657 | 6.21E-06 |
| *TTLL6* | BF | 12 | 24987347 | 25022530 | 0.582 | 6.47E-08 |
| *CSE1L* | BF | 17 | 50744910 | 50797548 | 0.729 | 3.49E-06 |
| *SYT12* | MS | 2 | 5410627 | 5433926 | 0.509 | 8.08E-06 |
| *CDKAL1* | MS | 7 | 15910666 | 16626276 | 0.785 | 1.83E-08 |
| *RAD51B* | MS | 7 | 91592424 | 92345294 | 0.989 | 2.52E-07 |
| *ABCD4* | MS | 7 | 97568208 | 97585655 | 0.852 | 3.82E-06 |
| *STIM2* | MS | 8 | 20517587 | 20604319 | 0.883 | 4.38E-07 |
| *ABL1* | WT | 1 | 270761668 | 270906708 | 0.937 | 7.35E-09 |
| *ST6GALNAC3* | WT | 6 | 136633441 | 137201252 | 0.980 | 1.12E-06 |
| *APBB2* | WT | 8 | 31921593 | 32193940 | 0.992 | 2.04E-06 |
| *RFTN1* | WT | 13 | 3454454 | 3674998 | 0.582 | 9.69E-06 |
| *RAB28* | NBA | 8 | 8774790 | 9125240 | 0.879 | 9.42E-06 |
| *ARHGEF38* | NBA | 8 | 116062408 | 116199137 | 0.963 | 7.41E-06 |
| *SMURF1* | NBD | 3 | 6037741 | 6142575 | 0.723 | 9.69E-06 |

Note: Gene boundaries were defined using Ensembl annotations (Sscrofa11.1) and extended by 3 kb upstream and downstream to include potential regulatory regions. Trait abbreviations: WT, off-test body weight; BF, back fat thickness; MS, loin muscle depth; NBA, number of piglets born alive; NBD, number of piglets born dead; NW, number of piglets weaned.

Table S6. AUPRC comparison between different GRM constructions.

| Sample size | Method | Number of causal variants | | | | | | Average |
|---|---|---|---|---|---|---|---|---|
| | | 1 | | 2 | | 3 | | |
| | | PVE=0.01 | PVE=0.04 | PVE=0.01 | PVE=0.04 | PVE=0.01 | PVE=0.04 | |
| 5,000 | FINEMAP-adj (Simulation-10k-SNP GRM) | 0.126 | 0.381 | 0.051 | 0.173 | 0.03 | 0.08 | 0.14 |
| | FINEMAP-adj (Random-10k-variant GRM) | 0.128 | 0.381 | 0.05 | 0.167 | 0.026 | 0.087 | 0.14 |
| | SuSiE-adj (Simulation-10k-SNP GRM) | 0.117 | 0.388 | 0.036 | 0.159 | 0.026 | 0.066 | 0.132 |
| | SuSiE-adj ((Random-10k-variant GRM) | 0.112 | 0.382 | 0.034 | 0.127 | 0.022 | 0.064 | 0.124 |
| 10,000 | FINEMAP-adj (Simulation-10k-SNP GRM) | 0.189 | 0.463 | 0.089 | 0.303 | 0.076 | 0.188 | 0.218 |
| | FINEMAP-adj (Random-10k-variant GRM) | 0.183 | 0.437 | 0.074 | 0.285 | 0.065 | 0.171 | 0.202 |
| | SuSiE-adj (Simulation-10k-SNP GRM) | 0.207 | 0.478 | 0.086 | 0.225 | 0.066 | 0.149 | 0.202 |
| | SuSiE-adj (Random-10k-variant GRM) | 0.192 | 0.488 | 0.08 | 0.215 | 0.062 | 0.144 | 0.197 |

Note: AUPRC values were calculated by varying PIP thresholds from 0 to 1 and computing the area under the resulting precision-recall curves. "Simulation-10k-SNP GRM" represents analyses using the same 10,000 chip SNPs used to construct the GRM during trait simulation, while "Random-10k-variant GRM" represents analyses using a different set of 10,000 randomly selected variants from imputed sequence data. Both approaches used GCTA-MLMA (LMM) summary statistics. Results are shown for different sample sizes (5,000 and 10,000 individuals), PVE levels (1% and 4%), and numbers of causal variants (1-3).
